# Host- and age-dependent transcriptional changes in *Mycobacterium tuberculosis* cell envelope biosynthesis genes after exposure to human alveolar lining fluid

**DOI:** 10.1101/2021.09.08.459334

**Authors:** Anna Allué-Guardia, Andreu Garcia-Vilanova, Angélica M. Olmo-Fontánez, Jay Peters, Diego J. Maselli, Yufeng Wang, Joanne Turner, Larry S. Schlesinger, Jordi B. Torrelles

## Abstract

Tuberculosis (TB) infection, caused by the airborne pathogen *Mycobacterium tuberculosis* (*M*.*tb*), resulted in almost 1.4 million deaths in 2019 and the number of deaths is predicted to increase by 20% over the next 5 years due to the COVID-19 pandemic. Upon reaching the alveolar space, *M*.*tb* comes in close contact with the lung mucosa before and after its encounter with host alveolar compartment cells. Our previous studies show that homeostatic innate soluble components of the alveolar lining fluid (ALF) can quickly alter the cell envelope surface of *M*.*tb* upon contact, defining subsequent *M*.*tb*-host cell interactions and infection outcomes *in vitro* and *in vivo*. We also demonstrated that ALF from 60+ year old elders (E-ALF) *vs*. healthy 18- to 45-year-old adults (A-ALF) is dysfunctional with loss of homeostatic capacity and impaired innate soluble responses linked to high local oxidative stress. In this study, a targeted transcriptional assay demonstrates that *M*.*tb* exposure to human ALF alters the expression of its cell envelope genes. Specifically, our results indicate that A-ALF-exposed *M*.*tb* upregulates cell envelope genes associated with lipid, carbohydrate, and amino acid metabolism, as well as genes associated with redox homeostasis and transcriptional regulators. Conversely, *M*.*tb* exposure to E-ALF shows lesser transcriptional response, with most of the *M*.*tb* genes unchanged or downregulated. Overall, this study indicates that *M*.*tb* responds and adapts to the lung alveolar environment upon contact, and that the host ALF status determined by factors such as age might play an important role in determining infection outcome.

## INTRODUCTION

*Mycobacterium tuberculosis* (*M*.*tb*), the causative agent of tuberculosis (TB), is one of the top leading causes of mortality worldwide due to a single infectious agent, with ∼1.4 million attributed deaths in 2019 [1]. However, global estimates indicate that worldwide disruptions in the healthcare system during the current COVID-19 pandemic could lead to an additional 6.3 million new TB cases between 2020 and 2025 and an added 1.4 million more TB deaths [2, 3]. Strict lockdowns have prevented patients from having access to TB medications and clinical evaluations, and have led to decreased TB diagnosis rates, since available resources have been redirected to prevent the spread of COVID-19 [4, 5]. These factors are predicted to cause not only an increase in the number of TB cases, but also to promote the development of drug-resistant TB, stressing the need for the development of new anti-TB therapies [6].

Most current drugs target the *M*.*tb* cell envelope, a highly complex and dynamic structure comprised mainly of carbohydrates and lipids, which provide structural support and resistance to osmotic changes, as well as a critical immunoregulatory role during *M*.*tb* infection [7-10]. It consists of four main layers: 1] an inner plasma membrane with periplasmic space; 2] a peptidoglycan (PG) core covalently linked to arabinogalactan (AG) and mycolic acids (MAcs); 3] a peripheral layer of non-covalently linked lipids, glycolipids, and lipoglycans [*e*.*g*. phthiocerol dimycocerosates (PDIMs), trehalose dimycolate (TDM) and monomycolate (TMM), sulfolipids (SLs), phosphatidyl-*myo*-inositol mannosides (PIMs), lipomannan (LM), and mannose-capped lipoarabinomannan (ManLAM), among others], and; 4) the outermost layer or capsule [11, 12]. The pathogenesis of *M*.*tb* is inherently linked to its heterogeneous and dynamic cell envelope surface, and cell envelope remodeling has been observed during infection and in response to environmental stresses [10]. Thus, the particular cell envelope composition of a mycobacterial cell at a given moment will define *M*.*tb*-host cell interactions and determine the infection outcome. However, it remains poorly understood how the *M*.*tb* cell envelope changes and adapts to the host lung environment during the natural course of pulmonary infection, a critical gap in our knowledge for defining new drug targets against *M*.*tb* relevant to bacteria in the lung environment.

After the host inhales droplets containing *M*.*tb*, the airborne pathogen is deposited in the lung alveolar space. Here it first comes in contact with soluble components of the alveolar lining fluid (ALF) for an undefined period of time (from minutes to hours/days) [13-16], before and after its encounter with host alveolar compartment cells such as alveolar macrophages (AMs) or alveolar epithelial cells (ATs), and immune cells such as neutrophils [17, 18]. Our previous studies have demonstrated that homeostatic ALF hydrolytic enzymes, whose function is to promote lung health, can modify the *M*.*tb* cell envelope without reducing *M*.*tb* viability [19, 20], including a reduction in major *M*.*tb* virulent factors mannose-capped lipoarabinomannan (ManLAM) and trehalose dimycolate (TDM), from the *M*.*tb* cell surface. These ALF-derived *M*.*tb* cell envelope modifications have an impact on *M*.*tb* infection outcomes *in vitro* and *in vivo*, since they allow for a better recognition by cells of the immune system and improved control of the infection [19-23]. Indeed, exposure to ALF results in decreased *M*.*tb* association and intracellular growth within human macrophages, as well as altered intracellular trafficking and increased pro-inflammatory responses [19]. Neutrophils also possess an enhanced innate ability to recognize and kill intracellular ALF-exposed *M*.*tb*, while limiting excessive inflammatory responses [21]. *In vivo* infections using ALF-exposed *M*.*tb* demonstrate better control of infection in the mouse model [22]. Further, *M*.*tb* fragments released after ALF exposure by the action of the ALF hydrolases, are capable of priming neutrophils and modulating macrophages in an IL-10 dependent manner to contribute further to the control of *M*.*tb* [20, 23].

Importantly, the levels and functionality of ALF soluble components are altered in certain human populations such as the elderly, with increased pro-oxidation and pro-inflammatory pathways, altered complement and surfactant levels, and decreased binding capability of surfactant protein A (SP-A) and D (SP-D) in the aging lung, defining what we call ‘dysfunctional ALF’. Consequently, *M*.*tb* exposed to elderly dysfunctional ALF, while maintaining its viability, show increased intracellular growth in macrophages and ATs, as well as increased bacterial burden in mice with increased lung tissue damage [22, 24, 25]. Altogether, these results indicate that the host ALF functional status plays a key role in shaping the *M*.*tb* cell envelope during the initial stages of infection. However, the overall impact of these different ALF microenvironments on *M*.*tb* adaptation to the human host and subsequent infection progression is still largely unknown.

Since hydrolytic enzymes present in functional human ALF (from healthy adult individuals) modify the *M*.*tb* cell envelope [19], we hypothesize that *M*.*tb* will compensate for these ALF-driven modifications by altering the expression of genes related to cell envelope biogenesis. Conversely, *M*.*tb* exposed to ALF with decreased functionality (such as elderly ALF) will show little to no changes. In addition, our published data [20, 26] demonstrate that 15-minute exposure to human ALF is enough to alter the *M*.*tb* cell envelope and its interactions with host cells and that these modifications are maintained for up to 24 h [20, 26]. In this study, we first aim to determine if short (15 min) or long (12 h) exposure to human ALF has any effects on the expression of targeted *M*.*tb* cell envelope genes, showing that gene expression is moderately altered at 15 min and that some transient expression changes happen at 12 h. Then, we use a multiplex qPCR assay to compare an extensive transcriptional profile of *M*.*tb* cell envelope genes associated with lipid, carbohydrate and amino acid metabolism, among others, after exposure to functional healthy adult (A)-*vs*. dysfunctional healthy elderly (E)-ALF. Our results show significant differences in gene expression, where A-ALF exposed *M*.*tb* upregulated genes involved in cell envelope remodeling, thus implicating this remodeling in subsequent *M*.*tb*-host interactions during the infection process.

## MATERIALS AND METHODS

### Human subjects and ethics statement

Human ALFs used in this study were previously isolated from collected bronchoalveolar lavage fluid (BALF) from healthy adult and elderly volunteers, in strict accordance with the US Code of Federal and approved Local Regulations (The Ohio State University Human Subjects IRB numbers 2012H0135 & 2008H0119 and Texas Biomedical Research Institute/UT-Health San Antonio/South Texas Veterans Health Care System Human Subjects IRB numbers HSC20170667H & HSC20170673H), and Good Clinical Practice as approved by the National Institutes of Health (NIAID/DMID branch), with written informed consent from all human subjects. Healthy adults (18-45 years old) and elderly (+60 years) were recruited from both sexes and no discrimination of race or ethnicity.

### Collection of BALF and ALF

BALF was collected from healthy adult or elderly donors in sterile endotoxin free 0.9% of NaCl, filtered through 0.2 μm filters, and further concentrated 20-fold using Amicon Ultra Centrifugal Filter Units with a 10-kDa molecular mass cut-off membrane (Millipore Sigma) at 4°C to obtain ALF with a physiological concentration reported within the human lung (1 mg/mL of phospholipid), as we previously described [19-24, 27]. ALF was aliquoted in low-protein binding tubes and stored at -80°C until further use.

### Bacterial cultures and ALF exposure

*M*.*tb* strains GFP-Erdman (kindly provided by Dr. Marcus Horwitz, UCLA) and H_37_R_v_ (ATCC# 25618) were cultured in 7H11 agar plates (BD BBL), supplemented with oleic acid, albumin, dextrose and catalase (OADC) at 37°C for 14 days. Single bacterial suspensions (∼1×10^9^ bacteria/ml) were obtained as previously described [19-23]. Bacterial pellets were exposed to individual ALFs (from different donors) for 15 min or ∼12 h at 37°C. After exposure, ALF was removed and bacterial pellets were directly incubated in RNAProtect (Qiagen) for 10 min at room temperature (RT), centrifuged at 13,000 x *g* and stored at -80°C until further use. For each of the ALFs, corresponding heat-inactivated ALFs (2 h at 80°C) [21] were used as controls in parallel.

### RNA extraction and cDNA synthesis

RNA from ALF-exposed bacterial pellets was extracted using the Quick-RNA Fungal/Bacterial Miniprep kit (Zymo Research), following the manufacturer’s protocol. Briefly, bacterial pellets were resuspended in Lysis buffer and transferred to a ZR BashingBead Lysis tube containing 0.1 mm and 0.5 mm ceramic beads. Bead beating procedure was performed to break the tough-to-lyse mycobacterial cell envelope in a Disruptor Genie [10 cycles of 1 min at maximum speed with 1 min intervals on ice]. RNA was isolated from the supernatant using Zymo-spin columns, including an in-column DNAse I treatment, and eluted in nuclease-free water. To completely remove the genomic DNA, a second DNAse treatment was performed on the isolated RNA using TURBO DNAse reagent (Thermo Fisher Scientific) for 30 min at 37°C. Final RNA concentration and quality were measured with a Qubit 4 Fluorometer using the HS RNA kit (Thermo Fisher Scientific), and a Nanodrop One^C^, respectively. RNA (500 ng) was used for the synthesis of cDNA using the RevertAid H Minus First Strand cDNA Synthesis kit (Thermo Fisher Scientific) with random hexamer primers, following the manufacturer’s guidelines.

### qPCR analysis of targeted genes

Real-time quantitative PCR (qPCR) was performed to measure the expression of ten genes associated with the *M*.*tb* cell wall biosynthesis pathways (*pmmA, manB, whiB2, pimA, pimB, pimF, embC, pmmB, pmm1, manA*) after exposure of *M*.*tb* H_37_R_v_ to healthy human ALF (n=2 biological replicates, from two different donors). cDNA and primers were used in a 20 μl qPCR reaction with PowerUp SYBR Green Master Mix (Applied Biosystems), following manufacturer’s instructions. Reactions were run in an Applied Biosystems 7500 Real-Time PCR instrument with the following settings: reporter SYBR Green, no quencher, passive reference dye ROX, standard ramp speed, and continuous melt curve ramp increment. Expression was calculated relative to housekeeping genes *rpoB* or *sigA* using the 2^−ΔΔCT^ method [28].

### High-throughput multiplex qPCR

Primer pairs for multiplex qPCR assay were designed using BatchPrimer3 (https://wheat.pw.usda.gov/demos/BatchPrimer3/) [29] and PrimerQuest (IDT, https://www.idtdna.com/pages/tools/primerquest), with the following settings: primer size 18-21 ntds, T_m_ of ∼ 54-60°C, and maximum 3’ self-complementarity of 3 ntd. Best primers within these parameters were selected and aligned to the *M*.*tb* H_37_R_v_ reference genome (Genbank accession number: NC_000962.3) [30-32] to confirm that they uniquely aligned to the targeted gene regions. A high-throughput multiplex qPCR targeting more than 80 genes associated with *M*.*tb* metabolism and cell wall biogenesis was performed in *M*.*tb* Erdman exposed to ALF using the Biomark 96.96 Dynamic Array IFC for Gene Expression in a Biomark HD instrument (Fluidigm). Briefly, 1.25 μl of cDNA was pre-amplified in a 5 μl reaction with our pool of specific primers (500nM) using the Fluidigm Preamp Master Mix for a total of 10 cycles. Then, a 1/10 dilution of the pre-amplified cDNA and 100 μM of combined forward and reverse primers were used to prepare the sample pre-mix and assay mix, respectively. The 96.96 IFC chip was loaded and the assay run in a Biomark HD following the manufacturer’s instructions for Gene expression using Delta Gene Assays. Relative expression for each of the genes was calculated in the Fluidigm Real-Time PCR Analysis software using the 2^−ΔΔCT^ method with *rpoB* as the reference gene, and reported as fold changes of A-ALF (n=3 biological replicates using ALFs from different donors) or E-ALF (n=3 biological replicates using ALFs from different donors) exposed *M*.*tb* Erdman compared to control samples (*M*.*tb* Erdman exposed to corresponding heat-inactivated ALFs). As a control to compare both methods (Biomark *vs*. targeted qPCR), we included *M*.*tb* H_37_R_v_ exposed to adult ALF in the Biomark multiplex qPCR assay.

### Statistical analysis

Statistical significance between the two qPCR methods used in this study was calculated in GraphPad Prism v9.0.1 for each of the genes with a two-way ANOVA using the Sidak’s correction for multiple comparisons test with a 95% confidence interval.

## RESULTS

### Exposure to human ALF alters the expression of cell envelope PIMs/LM/ManLAM biosynthesis genes in *M*.*tb*

Our laboratory has previously demonstrated that ALF hydrolases modify the *M*.*tb* cell envelope, and that these cell envelope modifications occur in as little as 15 min of *M*.*tb* being in contact with ALF and are maintained up to 24 h, without compromising *M*.*tb* viability [19]. Importantly, these ALF-driven alterations of the *M*.*tb* cell envelope had an effect on *M*.*tb* infection outcomes *in vitro* and *in vivo* [19-23]. Based on these results, we sought to determine if *M*.*tb* alters its cell wall biosynthetic pathways as a direct consequence of ALF exposure. Because one of the main *M*.*tb* cell envelope components, ManLAM, is decreased by the action of ALF hydrolases [19] and PIMs, LM and ManLAM are thought to be part of the same biosynthetic pathway, we first selected a few key *M*.*tb* cell envelope genes associated with biosynthesis of initial mannose donors GDP-Man [*manA/Rv3255c, manB/Rv3264c* (previously annotated as *manC*), *pmmA/ Rv3257c* (previously annotated as *manB*), *pmmB/ Rv3308*] and polyprenylphosphate-based mannose or PPM (*pmm1/Rv2051c*), the transcriptional regulator *whiB2/Rv3260c* [33], and mannosyl- (*pimA/Rv2610c, pimB/Rv2188c, pimF/ Rv1500*) and arabynosyl- (*embC/Rv3793*) transferases (**Fig 1** and **Table 1**) [34, 35], all involved in this pathway. Expression was determined by RT-qPCR from *M*.*tb* previously exposed to human ALF for 15 min and 12 h. Relative expression was normalized to housekeeping genes *rpoB* (**Fig 2A**) or *sigA* (**Fig 2B**). *M*.*tb* preparations exposed to the same heat-inactivated ALFs were used as reference samples.

**Fig 1.**
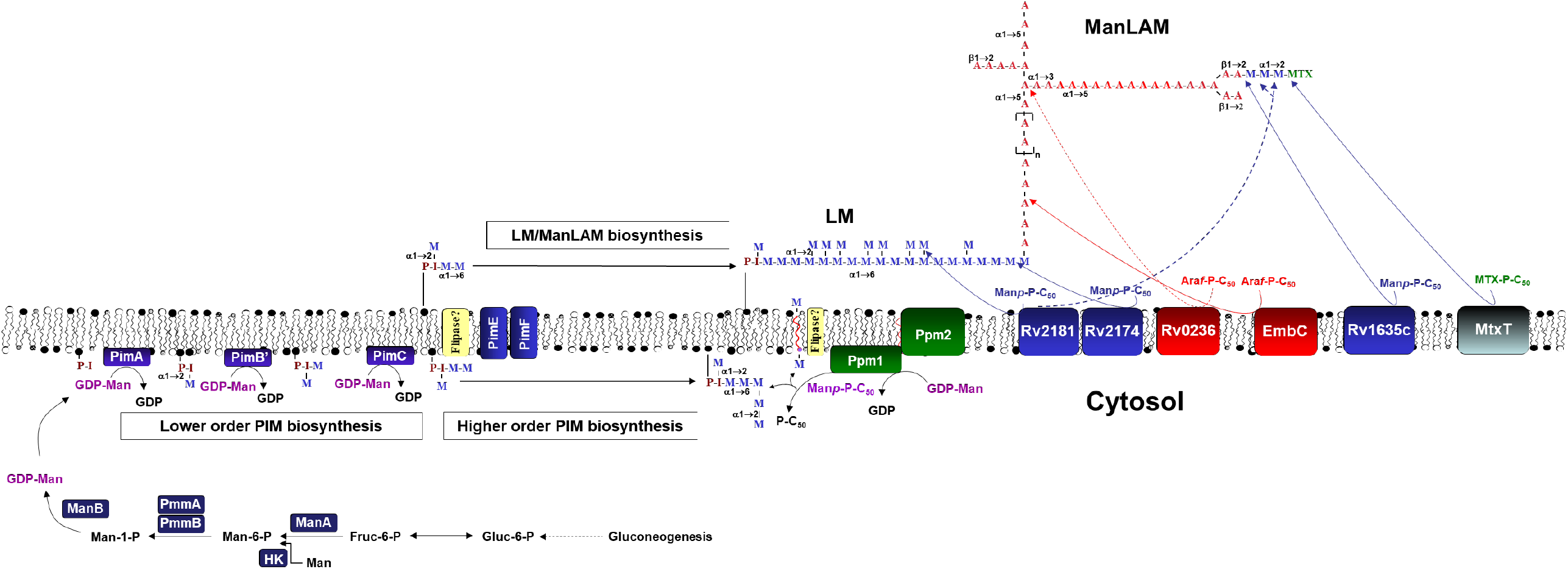
*M*.*tb* GDP-Man, PPM and PIM/LM/ManLAM biosynthetic pathways. GDP-Man can be biosynthesized directly from gluconeogenesis, through formation of Fruc-6-P from Glc-6-P (by the action of Glc-6-P isomerase), formation of Man-6-P (by the action of Man-6-P isomerase or ManA/Rv3255c), formation of Man-1-P (by the action of phosphomutases PmmA/Rv3257c, previously annotated as ManB, and PmmB/Rv3308), and finally formation of GDP-Man (by the action of Man-1-P guanylyn-transferase or ManB/Rv3264c, previously annotated as ManC). Further, Man-6-P can also be directly formed from Man by the action of a hexokinase (HK). PPM is formed from GDP-Man by the action of polyprenyl monophosphomannose synthase or Ppm1/Ppm2). Further, CDP-DAG together with inositol by the action of PI synthase (Rv2612c) forms PI, which is further mannosylated by several mannosyltransferases (PimA to PimF) to generate higher orders PIMs using GDP-Man and PPM as mannose donors. At one point, from PIM_4_ and using PPM as the major mannose donor, PIM_4_ is heavily mannosylated by an undisclosed number of mannosyl-transferases generating LM, and further arabinosylated with arabinosyl-transferases generating LAM, which can be further mannose-capped by the mannosyl-transferases action. LAM can also contain methylthio-_D_-xylose (MTX) capping motifs [78, 79], where MtxT (Rv0541c) transfers MTX to the mannoside caps of LAM [80]. Note: For simplicity, acyltransferases (*e*.*g*. Rv2610c) are not depicted, and neither is the formation of MTX-P-C50 by MtxS (Rv0539).

**Table 1.**
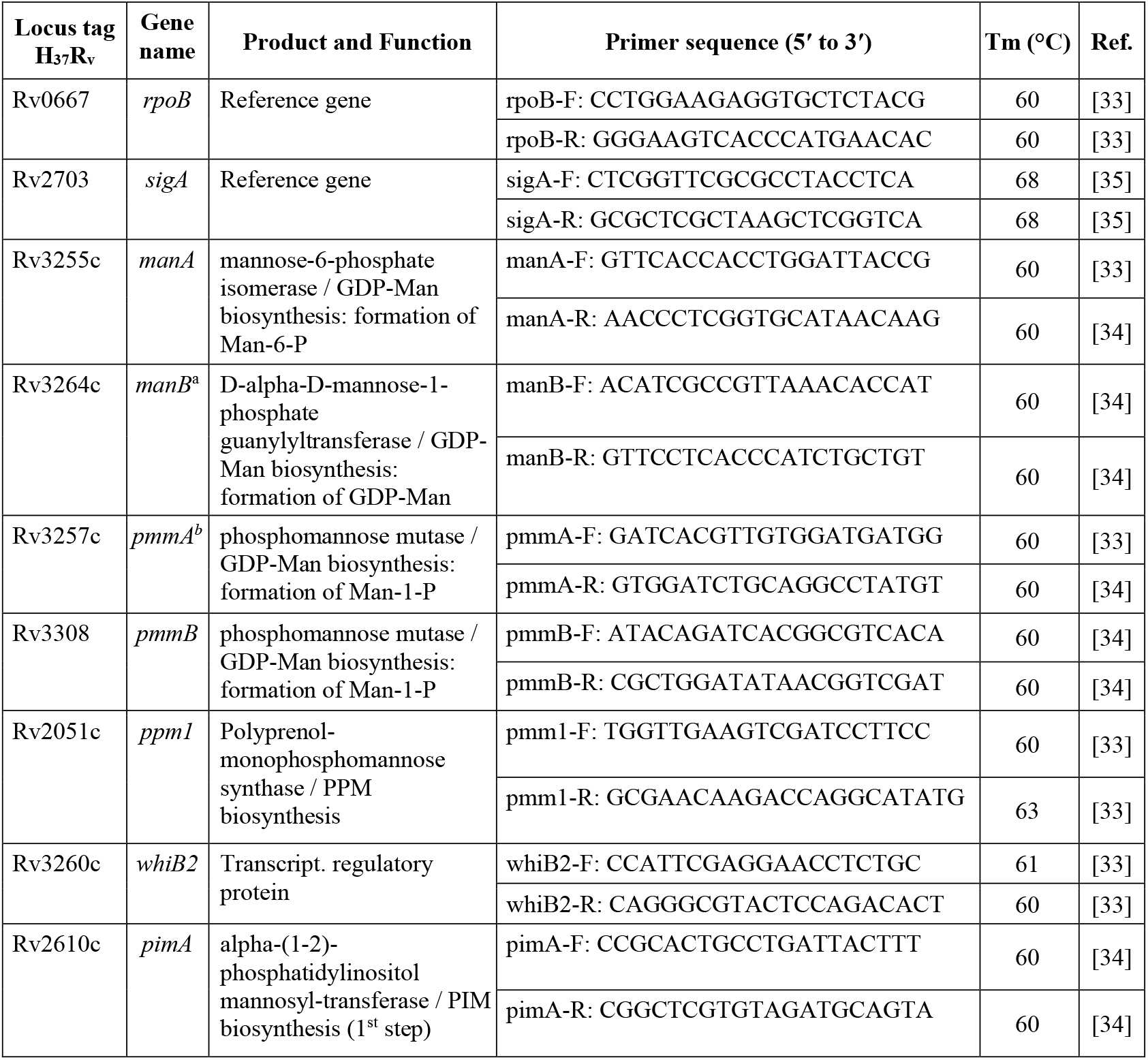
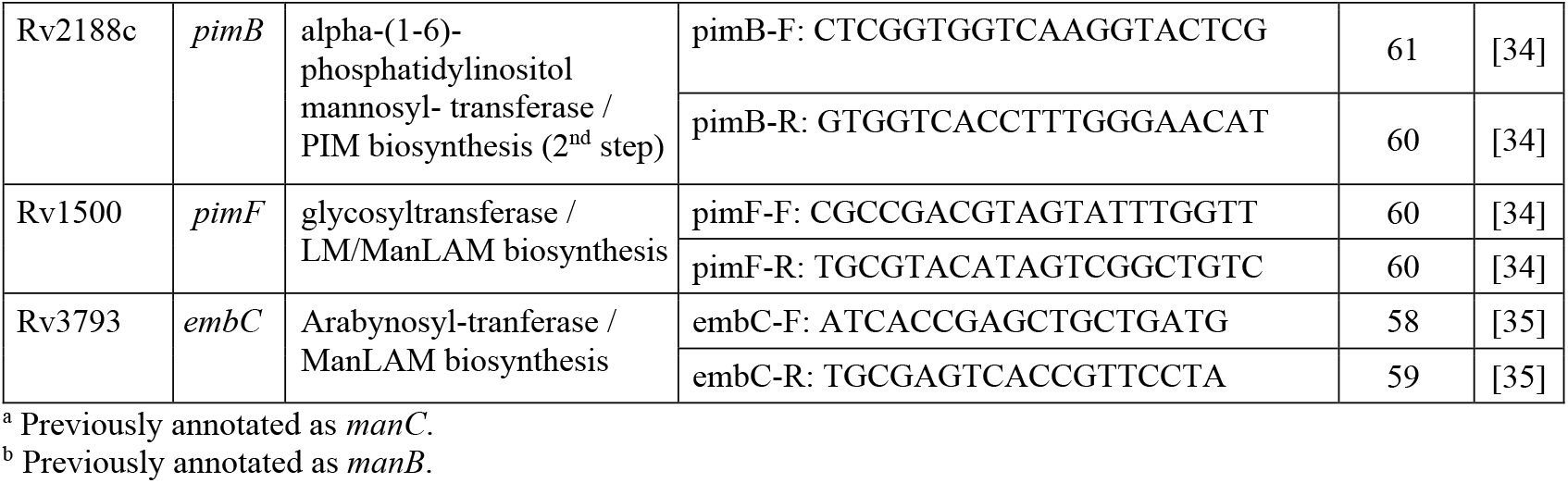
List of primers targeting the PIMs/LM/ManLAM biosynthesis pathways in *M*.*tb*.

**Fig 2.**
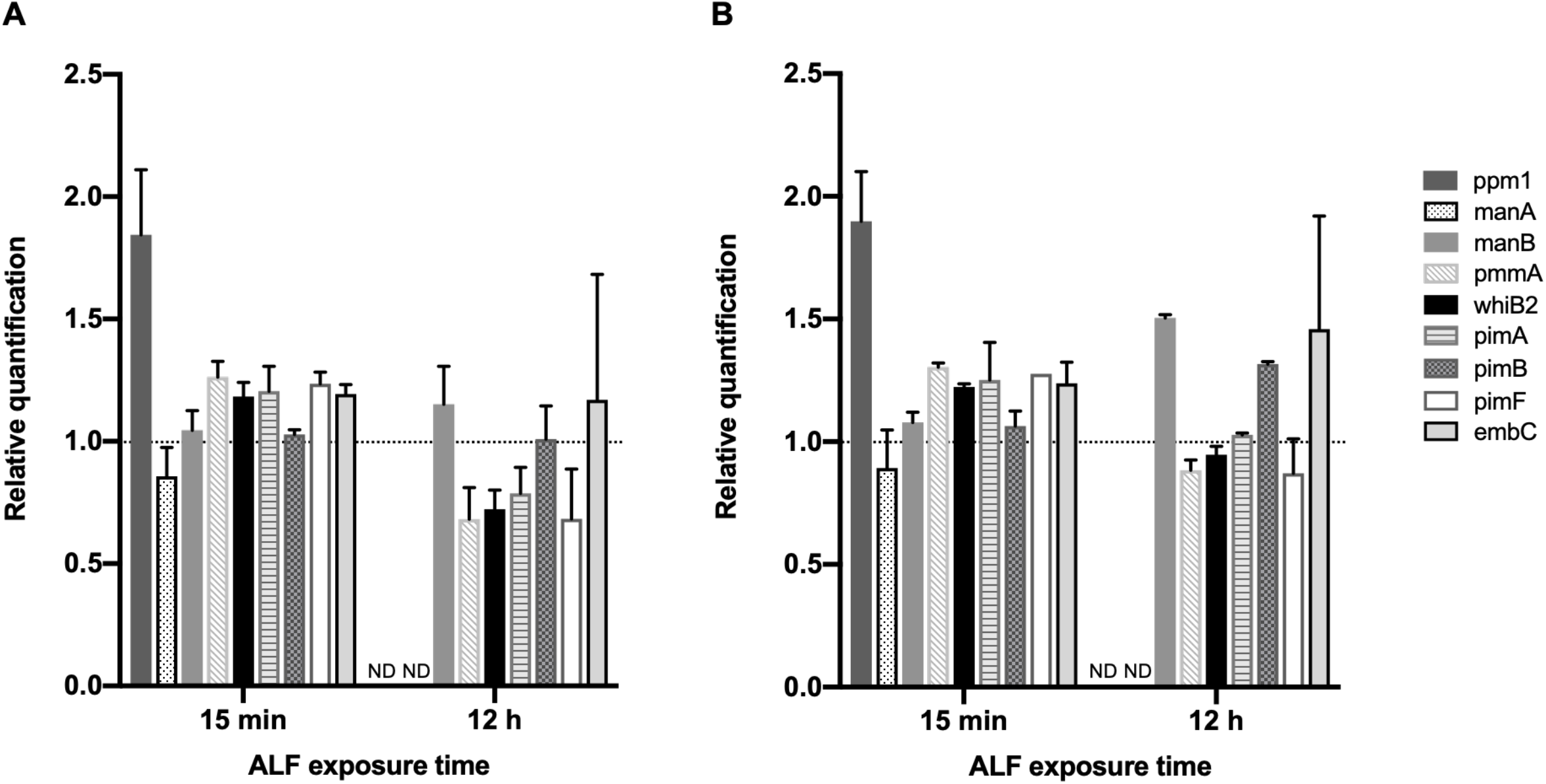
Relative expression of selected PIM/LM/ManLAM biosynthesis genes in *M*.*tb* H_37_R_v_ after exposure to ALF. *M*.*tb* was exposed for 15 min and 12 h to healthy human ALF (n=2 biological replicates, from two independent ALF donors), using *rpoB* (**A**) or *sigA* (**B**) as reference genes. Expression values are shown as fold changes, and were calculated using the 2^−ΔΔCT^ method (ALF-exposed *M*.*tb vs*. heat-inactivated ALF-exposed *M*.*tb*) and plotted as the mean ± SEM using Prism v9. ND: not detected (below limit of detection).

Several of the targeted *M*.*tb* cell envelope genes were moderately upregulated after 15 min of exposure to ALF when compared to *M*.*tb* exposed to heat-inactivated ALF (depicted as fold changes), except for *manA* that was slightly downregulated and *manB* and *pimB* that did not show any changes **(Fig 2)**. Indeed, *ppm1* (polyprenol-monophosphomannose synthase), which transfers mannose from GDP-mannose to endogenous PPM [36, 37] in the PIMs/LM/ManLAM biosynthesis pathway, showed the highest fold change (1.84 and 1.89 when using *rpoB* or *sigA* as reference genes, respectively) after 15 min of exposure compared to heat-inactivated ALF, although its expression was too low to be detected in our qPCR assay after 12 h (depicted as ND, **Fig 2**). Similarly, *manA* was below the detection limit at 12 h and expression differences could not be quantified (ND, **Fig 2**). A few of the tested genes showed some increased expression at 12 h post-ALF exposure (*manB, pimB*, and *embC*) compared to heat-inactivated ALF, while others showed no major transcriptional changes or were slightly downregulated.

When comparing 15 min *vs*. 12 h post-ALF exposure, some PIMs/LM/ManLAM biosynthesis-associated genes decreased their expression (*pmmA, whiB2, pimF, pimA*) while others increased (*manB, pimB, embC*) (**Fig 2** and **Table 2**). Although changes are only moderate, this suggests a temporal and dynamic adaptation of *M*.*tb*’s cell envelope components in response to ALF, in agreement with previous reports showing cell envelope remodeling of *M*.*tb* during infection and in response to different environmental conditions [33, 34, 38-43]. *pmmB* expression was too low to be detected by our qPCR assay at either timepoints tested, thus no fold changes are shown.

**Table 2.** Comparison of relative expression between 15 min and 12 h of ALF exposure for targeted qPCR genes. Differences in the expression of targeted *M*.*tb* cell envelope genes between 15 min and 12 h after ALF exposure were calculated (fold change at 12 h –fold change at 15 min) for each reference gene. Average differences for *rpoB* and *sigA* are also shown.

| Genes | 15 min vs. 12 h (*rpoB*) | 15 min vs. 12 h (*sigA*) | Mean *rpoB* and *sigA* |
| --- | --- | --- | --- |
| *manB* | 0.107 | 0.425 | 0.266 |
| *pmmA* | -0.580 | -0.421 | -0.500 |
| *whiB2* | -0.461 | -0.276 | -0.368 |
| *pimA* | -0.418 | -0.224 | -0.321 |
| *pimB* | -0.019 | 0.252 | 0.116 |
| *pimF* | -0.552 | -0.406 | -0.479 |
| *embC* | -0.024 | 0.222 | 0.099 |

### Effects of A-*vs*. E-ALF exposure on the expression of cell envelope biosynthesis genes in *M*.*tb*

Since we observed some changes in the expression of key *M*.*tb* cell envelope genes in response to ALF, and based on our previous publications showing that ALF status influences *M*.*tb*-host interactions via multiple factors, including age [22, 24], we next sought to determine if contact with different host ALFs would result in different transcriptional profiles of the *M*.*tb* cell envelope. We exposed *M*.*tb* Erdman to individual A-ALFs or E-ALFs for 12 h as described in our previous work [19], and calculated the relative expression of 83 genes related to cell envelope biosynthesis using a multiplex qPCR assay in a Biomark HD platform [44]. Results for some of the genes were compared to the previous targeted qPCR assay, showing concordance in gene expression between the two qPCR methods used in this study (**Supplemental Figure 1**). Small differences observed could be attributed to inherent variability within human ALF, since we used different human donors.

Genes selected for the multiplex qPCR are related to the biosynthesis of key *M*.*tb* cell envelope lipid and carbohydrate components (**Supplemental Table 1**). For lipid metabolism, we targeted genes from the following pathways: fatty acid metabolism, glycerolipid and glycerophospholipid metabolism, phosphatidylinositol (PI, precursor for more complex glycolipids such as PIM and LAM), PIMs/LM/ManLAM biosynthesis, mycolic acid biosynthesis, and linoleic acid metabolism [9]. For carbohydrate metabolism, we included genes related to: carbohydrate biosynthesis, glycolysis and gluconeogenesis, mannose and fructose metabolism, galactose metabolism, citrate cycle or TCA, glyoxylate and dicarboxylate metabolism, and inositol phosphate metabolism [45]. Gene names, with corresponding function, pathway and functional categories, as well as primer sequences are listed in **Supplemental Table 1**.

As shown in **Fig 3A**, most of the cell envelope genes associated with lipid metabolism from the different *M*.*tb-*targeted pathways were significantly upregulated in *M*.*tb* exposed to A-ALFs (A_1_ to A_3_, three different adult donors) when compared to the same *M*.*tb* strain exposed to E-ALFs (E_1_ to E_3_, three different elderly donors), with A_1_-ALF having the highest changes in gene expression (**Fig 3A** and **Supplemental Table 2**). Only a few genes were downregulated in *M*.*tb* exposed to A-ALFs, including *fabH* and *fadD25* (FA metabolism, the latter involved in lipid degradation), and *adhE1, glpK*, and *cdsA* (glycerolipid and glycerophospholipid synthesis). ATP-binding cassette transporter Rv1747, thought to be involved in the export of lipooligosaccharides (LOS) through the mycobacterial membrane, was also downregulated in A-ALF-exposed *M*.*tb*, whereas negative regulator of Rv1747, named Rv2623 [46], showed increased expression (**Fig. 3A**). Interestingly, *M*.*tb* exposed to two of the E-ALFs showed upregulation of Rv1747 (**Fig. 3A**), contrary to A-ALF exposure, indicating a potential increase in *M*.*tb* PIM export after exposure to elderly ALF. All genes involved in mycolic acid synthesis tested in this study were upregulated in *M*.*tb* exposed to A-ALF. Finally, exposure to E-ALFs showed lesser effects on the overall expression of *M*.*tb* lipid metabolism when compared to A-ALFs, with several genes showing moderate upregulation (especially in E_3_-ALF, which showed the highest upregulation among the elders for most of the genes), and others with no effects or even decreased expression (**Fig. 3A**).

**Fig 3.**
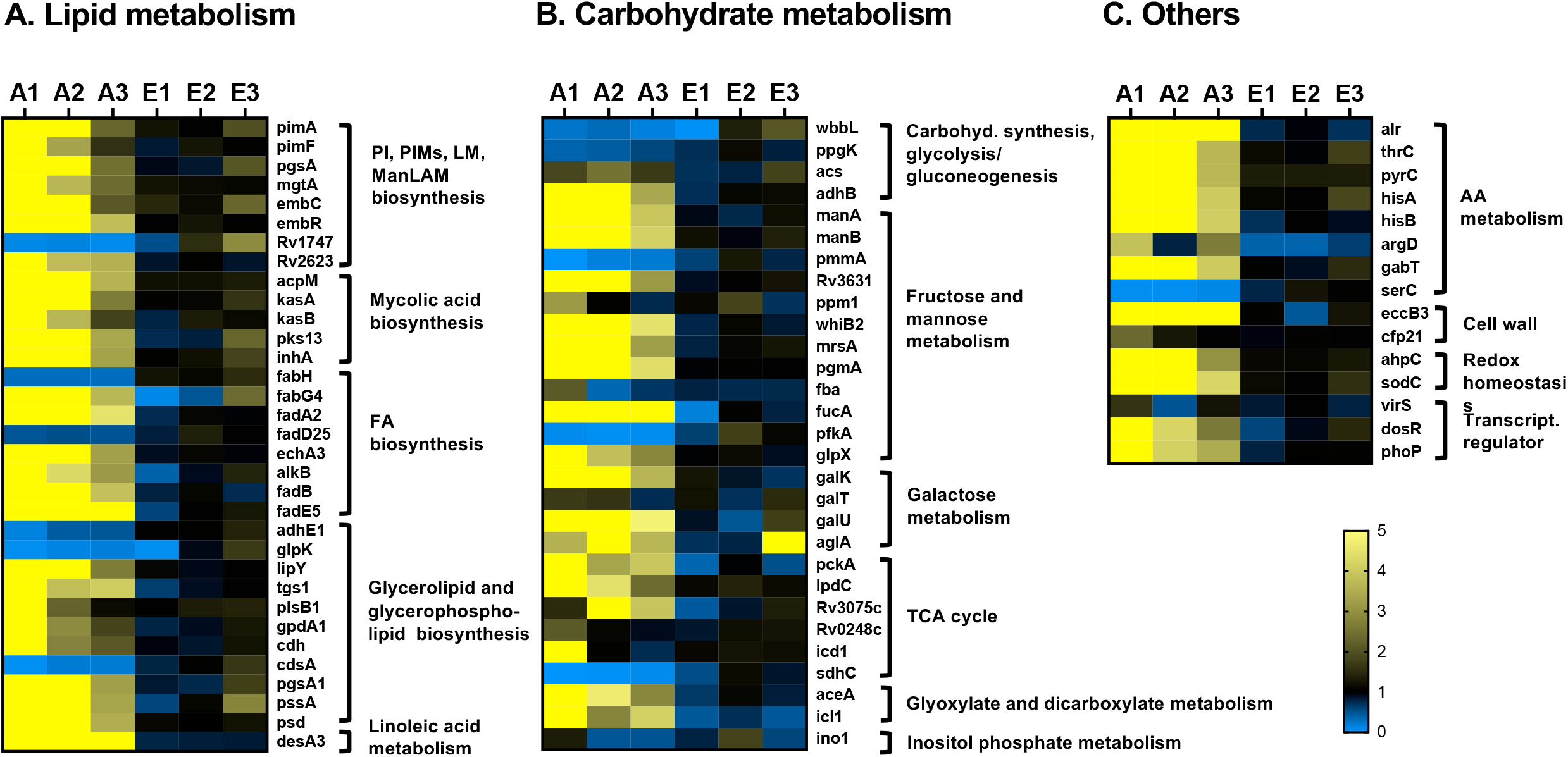
Relative expression of cell envelope biogenesis and metabolism genes in *M*.*tb* Erdman exposed to A-ALF and E-ALF. Heatmap showing relative expression of cell wall genes associated to (**A**) lipid metabolism; (**B**) carbohydrate metabolism; and (**C**) other pathways, in *M*.*tb* after being exposed to A-ALFs (n=3 biological replicates, A_1_- to A_3_-ALFs), or to E-ALFs (n=3 biological replicates, E_1_- to E_3_-ALFs). Expression was normalized using *rpoB* as reference gene and calculated using the 2^−ΔΔCT^ method (ALF-exposed *M*.*tb vs*. heat-inactivated ALF-exposed *M*.*tb*). Heatmap was constructed using Prism v9, with downregulated genes in blue (0 to 1 fold changes) and upregulated genes in yellow (1 to 5 or more fold changes). Genes are grouped based on their assigned pathways (see **Supplemental Table 1**). Note that genes *pimB, accD3, adhC* (lipid metabolism), *pmmB*, Rv0794c (carbohydrate metabolism), and *metZ* (others) were below the limit of detection in one or more of the samples, and have not been included in the heatmaps.

A similar trend was observed for carbohydrate metabolism genes (**Fig. 3B**), where most of the genes from the different pathways studied showed significant increased expression in *M*.*tb* exposed to A-ALFs when compared to E-ALFs. Particularly, 8 out of 12 genes associated with fructose metabolism and the mannose donor (GDP-Man/PPM) biosynthesis pathways were highly expressed in A-ALF-exposed *M*.*tb*, with only 2 genes downregulated (*pmmA*, which converts D-mannose 1-phosphate in D-mannose 6-phosphate, and *pfkA*, also a key enzyme involved in glycolysis) [47], and 2 other genes with variable expression across different A-ALF-exposed *M*.*tb* (*ppm1* and *fbA*) (**Fig 3B**). Galactose metabolism genes (*galK, galT, galU*, and *aglA*) essential for the biosynthesis of the cell envelope galactan core [48], were also upregulated after exposure to A-ALFs, while no changes or even decreased expression were observed in *M*.*tb* exposed to E-ALFs. Interestingly, predicted alpha-glucosidase *aglA* was significantly increased after exposure to E_3_-ALF (**Fig 3B**). Further, all tricarboxylic acid cycle (TCA)-associated genes were upregulated by A-ALF exposure, with the exception of *sdhC* (membrane-anchored subunit of Sdh2, implicated in *M*.*tb* growth linked to the TCA cycle under hypoxia conditions) [49] that was downregulated upon both A- and E-ALF *M*.*tb* contact. Glyoxylate and dicarboxylate genes were decreased in E-ALFs when compared to A-ALFs-exposed *M*.*tb*, whereas the only inositol phosphate gene tested *ino1*, catalyzing the first step in inositol synthesis for the production of major thiols and cell wall lipoglycans [50], had decreased expression after *M*.*tb* exposure to all ALFs tested, except for E_2_-ALF (**Fig 3B**).

In addition to lipid and carbohydrate pathways, we studied genes belonging to other categories such as amino acid metabolism, redox homeostasis, and transcriptional regulators (**Fig. 3C** and **Supplemental Table 1**). Results indicate that *M*.*tb* exposed to A-ALFs have major transcriptional changes compared to *M*.*tb* exposed to E-ALFs. Indeed, all genes associated with amino acid metabolism were highly upregulated in *M*.*tb* exposed to A-ALFs, except for *serC* (serine metabolism, **Fig 3C**) [51]. Other cell wall-associated genes were upregulated in *M*.*tb* exposed to A-ALFs including the putative membrane protein EccB3, part of the ESX-3 secretion system, important for zinc and iron uptake and homeostasis [52], and the cutinase precursor Cfp21, a lipolytic enzyme with immunogenic properties shown to elicit T and B cell responses [53, 54]. Similarly, genes *ahpC* and *sodC*, involved in the oxidative stress response [55, 56], and transcriptional regulators *dosR* and *phoP* had significantly increased expression in *M*.*tb* exposed to A-ALFs, whereas *virS* [57] was not significantly increased. Overall, our results demonstrate that exposure to functional A-ALF results in broad changes in *M*.*tb* cell envelope biosynthesis, suggesting highly dynamic cell envelope changes with constant remodeling within the lung alveolar environment, whereas *M*.*tb* exposed to dysfunctional E-ALF [22, 24, 25] showed more limited effects in gene expression.

## DISCUSSION

The cell envelope of *M*.*tb* is mainly composed of lipids and carbohydrates, and constitutes a dynamic structure known to adapt to the changing local host environment, especially during different stages of infection [8, 10]. Human ALF contains hydrolases whose homeostatic function is to maintain lung health. We have demonstrated that healthy adult individuals have up to 17 hydrolase activities capable of altering the *M*.*tb* cell wall. Indeed, exposure of *M*.*tb* to these adult ALFs significantly alters the *M*.*tb* cell wall reducing the content of two major cell envelope components, ManLAM and TDM, without compromising *M*.*tb* viability [19]. Importantly, these hydrolase activities are decreased in ALFs from healthy elders [24]. Thus, here, we demonstrate for the first time that exposure to human ALF, the first environment encountered by *M*.*tb* upon infection, results in transcriptional changes in key *M*.*tb* cell envelope biogenesis genes. Indeed, exposure to healthy A-ALFs resulted in increased expression of most of the *M*.*tb* cell envelope-associated genes from lipid, carbohydrate and amino acid metabolic pathways, among others. In contrast, *M*.*tb* exposed to E-ALFs from healthy elderly donors, which we demonstrated constitutes a more oxidized, pro-inflammatory and dysfunctional environment [22, 24], did not show many significant changes in gene expression.

We first assessed essential genes from the PIMs/LM/ManLAM synthesis and the mannose metabolism pathway (carbohydrate metabolism) (**Fig 1**), since our previous studies showed a decrease of ManLAM in the cell envelope of *M*.*tb* after ALF exposure [19]. PIM/LM/ManLAM molecules are essential to regulate *M*.*tb* recognition, uptake, survival and modulate the host immune response [58, 59]. Indeed, ManLAM has been shown to block phagosome-lysosome (P-L) fusion by inhibiting the Ca^2+^/Calmodulin phosphatidyl-inositol-3-kinase (PI3K) hvps34 pathway, promoting *M*.*tb* intracellular survival [60-62]. The biosynthesis of these molecules depends on mannose donors such as GPD-Man and PPM. Previous studies suggest that mannose donor levels are altered during the course of the infection. Indeed, *M*.*tb* mannose donor biosynthesis genes had increased expression levels 2 h after macrophage infection and then gradually decreased [33]. These were also found upregulated in an *in vitro* granuloma model [34].

In our study, most of the genes involved in the PIM/LM/ManLAM biosynthesis pathway were upregulated after exposure to A-ALFs (**Figs 3A** and **3B)**. We speculate that *M*.*tb* is trying to compensate for the loss of ManLAM and other mannose-containing cell envelope surface components due to the action of A-ALF hydrolases [19], with major implications for disease progression. In this regard, bacilli exposed to functional A-ALF might get taken up by antigen-presenting cells (APCs) before they can reconstitute ManLAM and will be cleared, while bacilli that rapidly upregulate and replenish ManLAM on the cell surface before its encounter with APCs will be able to block P-L fusion and survive withing the host cells. Conversely, exposure to E-ALFs did not have major transcriptional effects in genes from both mannose donors and PIM/LM/ManLAM biosynthesis pathways (**Figs 3A** and **3B)**, likely because E-ALF have less hydrolase activities [24]. This E-ALF deficiency in hydrolase activities could be directly linked to E-ALF being a highly oxidative environment [22, 24], therefore impacting the *M*.*tb* cell envelope and subsequent remodeling to a much lesser degree during infection. Since removal of surface lipids in *M*.*tb* enhances trafficking to acidic compartments [61], fewer ALF-driven alterations after E-ALF exposure might partially explain why *M*.*tb* replicates faster in the elderly lung environment by residing in its protective intracellular niche [22, 25]. Further studies will be necessary to determine the specific impact of elderly ALF hydrolases on the *M*.*tb* cell envelope.

Most lipid metabolism genes associated with mycolic acid synthesis, fatty acid metabolism, glycerolipid and glycerophospholipid synthesis, and linoleic acid metabolism were also significantly upregulated in A-ALF compared to E-ALF-exposed *M*.*tb* (**Fig 3A**). A similar trend was observed for carbohydrate metabolism, amino acid metabolism, and genes involved in redox homeostasis, stress response, and transcriptional regulation (**Figs 3B** and **3C**). Nitrogen and amino acid metabolism are important for *M*.*tb* pathogenesis and host colonization during infection, where intracellular bacteria exploit host nitrogen sources for growth and replication [63, 64]. Importantly, amino acids acquired from the host such as Ala and Gly might be directly assimilated for the synthesis of cell wall biomass and incorporated into the PG layer [65-67]. Transcriptional regulators DosR and PhoP also showed increased expression upon contact with functional A-ALF. These proteins are part of two-component systems implicated in a large number of *M*.*tb* adaptive responses, such as low oxygen levels during dormancy and persistence in the granuloma environment [68, 69], but they also play key roles as regulators of *M*.*tb* virulence [70, 71]. Interestingly, DosR has been shown to play a role in lipid accumulation during oxidative stress and iron starvation in certain *M*.*tb* clinical strains [72, 73].

These results indicate that the *M*.*tb* response to the ALF environment is not only limited to *M*.*tb* cell envelope remodeling, but also potentially affects overall *M*.*tb* metabolism and virulence [19-25, 27], with major implications in infection progression and TB disease outcome. Further global transcriptomic studies will be able to provide clues as to which other metabolic pathways are altered by the human ALF environment, with the potential to decipher key upregulated bacterial determinants during early stages of infection that can be targeted for the development of new preventative and therapeutic strategies [74]. Indeed, *M*.*tb* is exposed to ALF during the first infection stages upon deposition to the alveolar space, but also when escaping from necrotic cells or in cavities during active TB episodes leading to transmission [17]. Thus, it is plausible that *M*.*tb* adapts its cell envelope to the alveolar environment, upregulating the expression of specific genes to compensate the changes generated upon contact with ALF hydrolases and thus, determining interactions with host cells. In this regard, timing is expected to be important, as *M*.*tb* bacilli not able to restore its cell wall constitution before encountering antigen presenting cells such as alveolar macrophages might be cleared [19-21].

Key genes such as Rv1747, thought to participate in the export of PIMs to the cell envelope through negative regulation by stress protein Rv2623 [46], were highly downregulated in *M*.*tb* exposed to A-ALFs. Indeed, an ΔRv1747 mutant showed decreased levels of PIMs and was growth-attenuated, while ΔRv2623 had enhanced PIM expression and was hypervirulent in mice [75, 76]. Our data suggest that contact with functional A-ALF reduces the expression of PIM transporter Rv1747 through increased expression of Rv2623, modulating the export of immunomodulatory PIMs and potentially influencing bacterial growth and virulence.

Taken together, these results suggest that *M*.*tb* compensates for the loss of cell surface components (due to the action of ALF hydrolases) by upregulating and activating different cell envelope biosynthesis pathways to rebuild its cell wall, at the detriment of downregulating some key genes (*e*.*g*. Rv1747) involved in the transport of cell envelope components to the surface [19-21, 23, 46]. This shift between ALF innate homeostatic mechanisms and *M*.*tb* countermeasures in the ALF microenvironment, dependent on the host ALF status (A-AF *vs*. E-ALF), will likely determine subsequent interactions between *M*.*tb* and host cells, as well as intracellular trafficking and infection outcomes [19-21]. Further studies are needed to provide better insight to why E-ALF-exposed *M*.*tb* replicates faster than A-ALF-exposed *M*.*tb* in both professional and non-professional phagocytes, and to explain why E-ALF status in old age enhances *M*.*tb* infection *in vitro* and *in vivo* [22, 24, 25] and contributes to elders being more susceptible to respiratory infections in general. Finally, it is important to consider that the cell envelope composition of *M*.*tb* is strain-specific, with differences observed in *M*.*tb* strains from different lineages, and thus, different ALF-driven alterations in *M*.*tb* metabolism may drive different infection progression [77].

In summary, our study provides evidence that *M*.*tb* contact within the ALF shapes the composition of its cell envelope which, depending on the ALF status (‘functional A-ALF’ *vs*. ‘dysfunctional E-ALF’) [22, 24, 25], is likely to define subsequent *M*.*tb*-host cell interactions. Indeed, E-ALF-exposed *M*.*tb* presented minimal transcriptional changes when compared to A-ALF-exposed *M*.*tb*, which we speculate provides a fitness advantage to *M*.*tb* as its cell wall remains intact, and thus, it has the energy reserves required to efficiently infect and replicate faster within host cells of elderly individuals. In contrast, *M*.*tb* undergoes significant alterations on its cell wall (significant loss of virulent factors ManLAM and TDM, among others) upon exposure to A-ALF [19]. This triggers a greater transcriptional change that we interpret as efforts of A-ALF-exposed *M*.*tb* to reprogram its metabolism to quickly rebuild its cell wall, specifically upregulating genes involved in the biosynthesis of *M*.*tb* virulent factors such as ManLAM, with the ensuing energy requirements. This can be detrimental for A-ALF-exposed *M*.*tb* and favor host cells to control infection better in adult individuals. Future studies will investigate the metabolic status of E-ALF-exposed *M*.*tb* after infection of professional and non-professional phagocytes *in vitro* and *in vivo*, and will correlate bacterial and host determinants associated with increased susceptibility to infection in old age.

## AUTHORS CONTRIBUTIONS

AAG performed experiments and analyzed data. AGV analyzed data. AOF processed BAL samples. JP and DJM performed the bronchoalveolar lavage procedures in humans. AAG and JBT conceptually developed the study and wrote the manuscript. JT, LSS, and YW provided critical analysis of the data and editing of the manuscript. JT, LSS, and JBT provided funding. All authors read and approved the final version of this manuscript.

## ACKNOWLEDGEMENTS

We want to thank Sean Vargas from the University of Texas San Antonio (UTSA) Genomics Core for his assistance with the multiplex qPCR assay (Biomark HD).

