## Supplemental Material for "Host- and age-dependent transcriptional changes in *Mycobacterium tuberculosis* cell envelope biosynthesis genes after exposure to human alveolar lining fluid"

**Running Title:** Human lung mucosa and *M. tuberculosis* infection

**Keywords**: *Mycobacterium tuberculosis*, alveolar lining fluid (ALF), lung mucosa, cell envelope biosynthesis, gene expression

**SUPPLEMENTAL MATERIAL**

**TABLES**

**Supplemental Table 1. List of primers used for multiplex qPCR assay using the Biomark HD instrument.** The table includes gene name, locus tag, product, function, pathway and functional category obtained from Mycobrowser (https://mycobrowser.epfl.ch) and KEGG (https://www.kegg.jp), sequences for forward and reverse primers, primer melting temperature ($T_m$), 3' complementariety, and product size.

**Supplemental Table 2. Relative expression of *M.tb* cell envelope biogenesis genes.** Gene expression was obtained after multiplex qPCR assay with the Biomark 96.96 IFC Dynamic Array IFC for Gene Expression (Fluidigm). Values were calculated with the $2^{-\Delta\Delta CT}$ method (ALF-exposed *M.tb vs*. corresponding heat-inactivated ALF-exposed *M.tb*) using *rpoB* as the reference gene.

**FIGURES**

**Supplemental Fig 1. Comparison of gene expression results between targeted qPCR (qPCR) and multiplex qPCR (Biomark) for selected cell envelope genes in ALF-exposed *M.tb*.** *M.tb* was exposed to healthy human ALF (n=2 biological replicates for each of the assays). Relative expression was calculated with the $2^{-\Delta\Delta CT}$ method (ALF-exposed *M.tb vs.* heat-inactivated ALF-exposed *M.tb*) using *rpoB* as the reference gene. Values are plotted as the means ± SEM (Prism v9). Statistical significance between the two methods was calculated for each of the genes with a two-way ANOVA using the Sidak's correction for multiple comparisons test with a 95% confidence interval, ns: non-significant.

### Supplemental Table 1

| Category | Locus tag H37Rv | Gene name | Product | Function | Pathway | Functional Category (Mycobrowser) | Primer | Sequence | Tm (°C) | 3' compl | Product size | Reference |
|---|---|---|---|---|---|---|---|---|---|---|---|---|
| Reference gene | Rv0667 | **rpoB** | DNA-directed RNA polymerase (beta chain) RpoB (transcriptase beta chain) (RNA polymerase beta subunit) | Catalyzes the transcription of DNA into RNA using the four ribonucleoside triphosphates as substrates [catalytic activity: N nucleoside triphosphate = N diphosphate + {RNA}(N)] | RNA polymerase | information pathways | rpoB-F | ACGAGTGCAAAGACAAGG | 55.21 | 0 | 94 | This study |
| | | | | | | | rpoB-R | GTCTGACTCTTGATCTCACC | 54.92 | 0 | | This study |
| Lipid metabolism | Rv2610c | **pimA** | alpha-(1-2)-phosphatidylinositol mannosyltransferase PimA | Involved in the first mannosylation step in phosphatidylinositol mannoside biosynthesis (transfer of mannose residues onto PI, leading to the synthesis of phosphatidylinositol monomannoside) | PI, PIMs, LM and ManLAM biosynthesis | lipid metabolism | pimA-F | TGCCTGATTACTTTGTCTCC | 55.06 | 0 | 131 | This study |
| | | | | | | | pimA-R | TACGTCGAAATCACCATGC | 55.75 | 3 | | This study |
| Lipid metabolism | Rv2188c | **pimB** | alpha-(1-6)-phosphatidylinositol monomannoside | Involved in PIM biosynthesis | PI, PIMs, LM and ManLAM biosynthesis | lipid metabolism | pimB-F | GGTTTGCTTCTGCGTTCG | 57.48 | 2 | 151 | This study |
| | | | | | | | pimB-R | CGCGACAGACACACTACC | 57.85 | 0 | | This study |
| Lipid metabolism | Rv1500 | **pimF** | glycosyltransferase PimF | Function unknown | PI, PIMs, LM and ManLAM biosynthesis | intermediary metabolism and respiration | pimF-F | CGACTCAGACTTGGAAGAGG | 57.58 | 0 | 92 | This study |
| | | | | | | | pimF-R | CGTGGCAACCAAATACTACG | 59.11 | 2 | | This study |
| Lipid metabolism | Rv2612c | **pgsA** | PI synthase PgsA1 (phosphatidylinositol synthase) (CDP-diacylglycerol--inositol-3-phosphatidyltransferase) | Catalyzes the transfer of a free alcohol (inositol) onto CDP-diacylglycerol. The product of this putative ORF seems be essential to mycobacteria [catalytic activity: CDP-diacylglycerol + myo-inositol = CMP + phosphatidyl 1D-myo-inositol]. PgsA1 utilises inositol-phosphate rather than inositol, in contrast to its Eukaryotic ortholog (see Gräve et al. 2019) | PI, PIMs, LM and ManLAM biosynthesis | lipid metabolism | pgsA-F | TCTGCTGTGGTGGATAGC | 56.31 | 2 | 99 | This study |
| | | | | | | | pgsA-R | GCCTTGATGTAAGAGATCACC | 56.12 | 2 | | This study |
| Lipid metabolism | Rv0557 | **mgtA** | Mannosyltransferase MgtA, GDP-mannose-dependent alpha-mannosyltransferase similar to PimB | Involved in lipomannan (LM) biosynthesis | PI, PIMs, LM and ManLAM biosynthesis | lipid metabolism | mgtA-F | CACGGTATGGCAAAGAGC | 56.19 | 2 | 62 | This study |
| | | | | | | | mgtA-R | ACCGGAATGTACGAAGACG | 57.29 | 2 | | This study |
| Lipid metabolism | Rv3793 | **embC** | Integral membrane indolylacetylinositol arabinosyltransferase EmbC (arabinosylindolylacetylino sitol synthase) | Involved in the biosynthesis of the mycobacterial cell wall arabinan and resistance to ethambutol (EMB; dextro-2,2'-(ethylenediimino)-DI-1-butanol). Polymerizes arabinose into the arabinan of arabiogalactan [catalytic activity: UDP-L-arabinose + indol-3-ylacetyl-myo-inositol = UDP + indol-3-ylacetyl-myo-inositol L-arabinoside] | PI, PIMs, LM and ManLAM biosynthesis | cell wall and cell processes | embC-F | TCAAAGACGACTGGTTTAGG | 55.04 | 1 | 91 | This study |
| | | | | | | | embC-R | GTTCCTAGGTTCAGATCAGC | 55.26 | 2 | | This study |
| Lipid metabolism | Rv1267c | **embR** | transcriptional regulatory protein EmbR | Involved in transcriptional mechanism. Thought to regulate the biosynthesis of the mycobacterial cell wall arabinan and resistance to ethambutol (EMB; dextro-2,2'-(ethylenediimino)-DI-1-butanol), regulating EMBA\|Rv3794 and EMBB\|Rv3795 | Trancription factor: regulation of lipomannan/lipoarabinomannan ratio | regulatory proteins | embR-F | TCACCGCCTACTACCTCTCC | 57.1 | 0 | 74 | This study |
| | | | | | | | embR-R | GGCCAGTGTTGTCTTCACC | 55.9 | 1 | | This study |
| Lipid metabolism | Rv2623 | **Rv2623** | Universal stress protein family protein TB31.7n | Might negatively regulate transporter Rv1747 | Not assigned | virulence, detoxification, adaptation | Rv2623-F | GTGGTTGAACAGGCTTCGC | 59.72 | 2 | 135 | This study |
| | | | | | | | Rv2623-R | CCCACTTCCGAGACAACCC | 60 | 0 | | This study |
| Lipid metabolism | Rv1747 | **Rv1747** | conserved transmembrane ATP-binding protein ABC transporter | Thought to be involved in active transport of undetermined substrate (possibly lipooligosaccharide) across the membrane. Responsible for energy coupling to the transport system and for the translocation of the substrate across the membrane | Transporter | cell wall and cell processes | Rv1747-F | AGGTTACTTCGTCTTTCTGG | 54.73 | 0 | 139 | This study |
| | | | | | | | Rv1747-R | CAGCAACACTAGGATCTGG | 54.99 | 2 | | This study |
| Lipid [illegible] | Rv2244 | **acpM** | Meromycolate extension [illegible] | Involved in fatty acid biosynthesis [illegible] | mycolic acid biosynthesis | [illegible] | acpM-F | AGACCGAGGACAAGTACGG | 58.44 | 1 | [illegible] | This study |

| | | | | | | | | | | | | |
|---|---|---|---|---|---|---|---|---|---|---|---|---|
| metabolism | | | acyl carrier protein AcpM | (mycolic acids synthesis); involved in meromycolate extension | | lipid metabolism | acpM-R | TCGAGCTTCTGGATGTAGGC | 59.25 | 2 | 94 | This study |
| Lipid metabolism | Rv2245 | **kasA** | 3-oxoacyl-[acyl-carrier protein] synthase 1 KasA (beta-ketoacyl-ACP synthase) (KAS I) | Involved in fatty acid biosynthesis (mycolic acids synthesis); involved in meromycolate extension. Catalyzes the condensation reaction of fatty acid synthesis by the addition to an acyl acceptor of two carbons from malonyl-ACP [catalytic activity: acyl-[acyl-carrier protein] + malonyl-[acyl-carrier protein] = 3-oxoacyl-[acyl-carrier protein] + [acyl-carrier protein] + CO(2)] | mycolic acid biosynthesis | lipid metabolism | kasA-F | ACGTGGAAGGGTCTGTTGG | 56.8 | 0 | 76 | This study |
| | | | | | | | kasA-R | CTAGATCCCACTTGGTGACG | 54.5 | 2 | | This study |
| Lipid metabolism | Rv2246 | **kasB** | 3-oxoacyl-[acyl-carrier protein] synthase 2 KasB (beta-ketoacyl-ACP synthase) (KAS I) | Involved in fatty acid biosynthesis (mycolic acids synthesis); involved in meromycolate extension. Catalyzes the condensation reaction of fatty acid synthesis by the addition to an acyl acceptor of two carbons from malonyl-ACP [catalytic activity: acyl-[acyl-carrier protein] + malonyl-[acyl-carrier protein] = 3-oxoacyl-[acyl-carrier protein] + [acyl-carrier protein] + CO(2)] | mycolic acid biosynthesis | lipid metabolism | kasB-F | ATCTGGTAAACCTCGATCC | 53.62 | 2 | 88 | This study |
| | | | | | | | kasB-R | GAGTTATTGATCGCATACCG | 54.13 | 2 | | This study |
| Lipid metabolism | Rv3800c | **pks13** | Polyketide synthase Pks13 | Involved in the final steps of mycolic acid biosynthesis. Catalyses the condensation of two fatty acyl chains | mycolic acid biosynthesis | lipid metabolism | pks13-F | GCAACTGGGTGGGTAAGG | 55 | 0 | 126 | This study |
| | | | | | | | pks13-R | CGGTCAGGTCTTCTATGTCG | 54.7 | 2 | | This study |
| Lipid metabolism | Rv1484 | **inhA** | NADH-dependent enoyl-[acyl-carrier-protein] reductase InhA (NADH-dependent enoyl-ACP reductase) | This isozyme is involved in mycolic acid biosynthesis. Second reductive step in fatty acid biosynthesis. Involved in the resistance against the antituberculosis drugs isoniazid and ethionamide [catalytic activity: acyl-[acyl-carrier protein] + NAD(+) = trans-2,3-dehydroacyl-[acyl-carrier protein] + NADH] | mycolic acid biosynthesis, fatty acid biosynthesis | lipid metabolism | inhA-F | ATTCGATTGGGTTCATGCCG | 58.98 | 2 | 83 | This study |
| | | | | | | | inhA-R | GATGCCCTTGGACACATCC | 58.2 | 2 | | This study |
| Lipid metabolism | Rv0904c | **accD3** | acetyl-coenzyme A carboxylase carboxyl transferase (subunit beta) AccD3 (accase beta chain) | This protein is a component of the acetyl coenzyme A carboxylase complex; first, biotin carboxylase catalyzes the carboxylation of the carrier protein and then the transcarboxylase transfers the carboxyl group to form malonyl-CoA [catalytic activity ATP + acetyl-CoA + HCO(3)(-) = ADP + phosphate + malonyl-CoA] | fatty acid metabolism | lipid metabolism | accD3-F | GGTAGCCGACTCCTATGC | 56.61 | 2 | 132 | This study |
| | | | | | | | accD3-R | AGGAAGTCGAACTCACAGG | 56.4 | 0 | | This study |
| Lipid metabolism | Rv0533c | **fabH** | 3-oxoacyl-[acyl-carrier-protein] synthase III FabH (beta-ketoacyl-ACP synthase III) (KAS III) | Involved in fatty acid biosynthesis. Catalyzes the condensation reaction of fatty acid synthesis by the addition to an acyl acceptor of two carbons from malonyl-ACP. KAS III catalyzes the first condensation reaction which initiates fatty acid synthesis and may therefore play a role in governing the total rate of fatty acid production. Possesses both acetoacetyl-ACP synthase and acetyl transacylase activities [catalytic activity: acyl-[acyl-carrier protein] + malonyl-[acyl-carrier protein] = 3-oxoacyl-[acyl-carrier protein] + CO2 + [acyl-carrier protein]] | fatty acid metabolism | lipid metabolism | fabH-F | CCCACGATAGACATGTACG | 54.95 | 2 | 97 | This study |
| | | | | | | | fabH-R | GTCCAATGCCTTGAAACG | 54.42 | 2 | | This study |
| Lipid metabolism | Rv0242c | **fabG4** | 3-oxoacyl-[acyl-carrier protein] reductase FabG4 (3-ketoacyl-acyl carrier protein reductase) | Involved in the fatty acid biosynthesis pathway (first reduction step) [catalytic activity: (3R)-3-hydroxyacyl-[acyl-carrier protein] + NADP+ = 3-oxoacyl-[acyl-carrier protein] + NADPH] | fatty acid metabolism | lipid metabolism | fabG4-F | GACCTGGTAGGAAACAACC | 55.11 | 2 | 124 | This study |
| | | | | | | | fabG4-R | GTACCGGAGTAAAGAACTCG | 55.36 | 2 | | This study |
| Lipid metabolism | Rv0243 | **fadA2** | acetyl-CoA acyltransferase FadA2 (3-ketoacyl-CoA | Function unknown, but involved in lipid degradation [catalytic activity: acyl-CoA | fatty acid metabolism | lipid metabolism | fadA2-F | GGCATCAAACGCGTAGACC | 59.29 | 0 | 138 | This study |

| | | | | | | | | | | | | |
|---|---|---|---|---|---|---|---|---|---|---|---|---|
| [illegible] | | | [illegible] thiolase) (beta-ketothiolase) | [illegible] + acetyl-CoA = CoA + 3-oxoacyl-CoA] | | [illegible] | fadA2-R | GTTAGGCCGCAGATTGTCG | 58.99 | 2 | [illegible] | This study |
| Lipid metabolism | Rv0632c | **echA3** | enoyl-CoA hydratase EchA3 (enoyl hydrase) (unsaturated acyl-CoA hydratase) (crotonase) | Could possibly oxidize fatty acids using specific components [catalytic activity: (3S)-3-hydroxyacyl-CoA = trans-2(or 3)-enoyl-CoA + H(2)O] | fatty acid metabolism | lipid metabolism | echA3-F | GGCTTCGACCTGAAGATCC | 54.9 | 2 | 96 | This study |
| | | | | | | | echA3-R | GTAGGACAAGAGGCGATACG | 54.7 | 2 | | This study |
| Lipid metabolism | Rv1521 | **fadD25** | fatty-acid-AMP ligase FadD25 (fatty-acid-AMP synthetase) (fatty-acid-AMP synthase) | Function unknown, but involvement in lipid degradation | fatty acid metabolism | lipid metabolism | fadD25-F | ACAGCTAAGAGAACATGGG | 54.15 | 0 | 67 | This study |
| | | | | | | | fadD25-R | ATAGTCGAGGCTTTGTGG | 53.95 | 0 | | This study |
| Lipid metabolism | Rv3252c | **alkB** | transmembrane alkane 1-monooxygenase AlkB (alkane 1-hydroxylase) (lauric acid omega-hydroxylase) (omega-hydroxylase) (fatty acid omega-hydroxylase) (alkane hydroxylase-rubredoxin) | Thought to be involved in fatty acid metabolism. Generates octanol and oxidized rubredoxin from octane and reduced rubredoxin. Also hydroxylates fatty acids in the omega-position [catalytic activity: octane + reduced rubredoxin + (O)2 = 1-octanol + oxidized rubredoxin + H(2)O] | fatty acid metabolism | lipid metabolism | alkB-F | CAAACCTCAGTTGGCTCGG | 58.75 | 1 | 132 | This study |
| | | | | | | | alkB-R | GCGAATCCTTCTTGTGCCC | 58.29 | 2 | | This study |
| Lipid metabolism | Rv0860 | **fadB** | fatty oxidation protein FadB | Involved in fatty acid degradation (probably in fatty acid beta-oxidation cycle) | fatty acid metabolism, benzoate, caprolactam and butanoate metabolism | lipid metabolism | fadB-F | TCCAGTGGGACAAGGATGC | 59 | 2 | 74 | This study |
| | | | | | | | fadB-R | CATCACGTTGGTTGACCCG | 59 | 2 | | This study |
| Lipid metabolism | Rv0244c | **fadE5** | acyl-CoA dehydrogenase FadE5 | Function unknown, but involved in lipid degradation | fatty acid metabolism, propanoate metabolism | lipid metabolism | fadE5-F | ACCCAGATGACCGACAAGAC | 59.39 | 1 | 212 | Guirado et al., 2015 |
| | | | | | | | fadE5-R | ACTACCGGCAACATCAGGTC | 59.75 | 1 | | Guirado et al., 2015 |
| Lipid metabolism | Rv0162c | **adhE1** | zinc-type alcohol dehydrogenase (E subunit) AdhE1 | Dehydrogeneses a alcohol (OXIDO-reduction) [catalytic activity: an alcohol + NAD+ = an aldehyde or ketone + | fatty acid metabolism, glycerolipid metabolism | intermediary metabolism and respiration | adhE1-F | TGACGTATACAGACGTTCG | 54.56 | 2 | 95 | This study |
| | | | | | | | adhE1-R | CGGTGTGTAGATCTCATGG | 54.81 | 2 | | This study |
| Lipid metabolism | Rv3045 | **adhC** | NADP-dependent alcohol dehydrogenase AdhC | Generates aldehyde or ketone from alcohol [catalytic activity: alcohol + NADP(+) = aldehyde or ketone + NADPH] | glycerolipid metabolism | intermediary metabolism and respiration | adhC-F | AACCGAATTACCCTGTGG | 54.13 | 1 | 193 | This study |
| | | | | | | | adhC-R | GAGTTGTAGGTGAAGTTTGC | 54.13 | 2 | | This study |
| Lipid metabolism | Rv3696c | **glpK** | glycerol kinase GlpK (ATP:glycerol 3-phosphotransferase) (glycerokinase) (GK) | Acts in rate-limiting step in glycerol utilization. Key enzyme in the regulation of glycerol uptake and metabolism [catalytic activity: ATP + glycerol = ADP + glycerol 3-phosphate] | glycerolipid metabolism | intermediary metabolism and respiration | glpK-F | CGATTGTATGGCAGGATACC | 55.78 | 2 | 117 | This study |
| | | | | | | | glpK-R | CGCCAGAGAAATAAGTTGC | 54.52 | 2 | | This study |
| Lipid metabolism | Rv3097c | **lipY** | PE-PGRS family protein, triacylglycerol lipase LipY (esterase/lipase) (triglyceride lipase) (tributyrase) (triglyceride lipase) (tributyrase) | Generates diacylglycerol and a fatty acid anion from triacylglycerol [catalytic activity: triacylglycerol + H(2)O = diacylglycerol + a fatty acid anion] | glycerolipid metabolism | PE/PPE | lipY-F | GCTGTCGTGGTTTCTTGG | 56.35 | 0 | 84 | This study |
| | | | | | | | lipY-R | CCGTCATAGGTGGTGTACTGG | 59.86 | 0 | | This study |
| Lipid metabolism | Rv3130c | **tgs1** | Triacylglycerol synthase (diacylglycerol acyltransferase) Tgs1 | Involved in synthesis of triacylglycerol | glycerolipid metabolism, TAG biosynthesis | lipid metabolism | tgs1-F | GACCATGCAGTCGCAATCC | 59.28 | 1 | 126 | This study |
| | | | | | | | tgs1-R | ATCTCACTGGCACCCTTGG | 59.31 | 2 | | This study |
| Lipid metabolism | Rv1551 | **plsB1** | acyltransferase PlsB1 | Thought to be involved in lipid metabolism | glycerolipid and glycerophospholipid metabolism | lipid metabolism | plsB1-F | CCTGAACTTCTTTCCGATGG | 56.14 | 2 | 86 | This study |
| | | | | | | | plsB1-R | GGTAGACGGGAATATCTTTCG | 55.69 | 2 | | This study |
| Lipid metabolism | Rv0564c | **gpdA1** | glycerol-3-phosphate dehydrogenase [NAD(P)+] GpdA1 (NAD(P)H-dependent glycerol-3-phosphate dehydrogenase) (NAD(P)H-dependent dihydroxyacetone-phosphate reductase) | Involved in de novo phospholipid biosynthesis; glycerol-3 phosphate formation [catalytic activity: SN-glycerol 3-phosphate + NAD(P)+ = glycerone phosphate + NAD(P)H] | glycerophospholipid metabolism | lipid metabolism | gpdA1-F | GCTATCAGCAATGTTTCGC | 55.3 | 2 | 130 | This study |
| | | | | | | | gpdA1-R | GATACCCAGCGAATAGCC | 54.65 | 1 | | This study |
| Lipid metabolism | Rv2289 | **cdh** | CDP-diacylglycerol pyrophosphatase Cdh (CDP-diacylglycerol diphosphatase) (CDP-diacylglycerol phosphatidylhydrolase) | Involved in phospholipid biosynthesis [catalytic activity: CDP-diacylglycerol + H(2)O = CMP + phosphatidate] | glycerophospholipid metabolism | lipid metabolism | cdh-F | GGTCGCAATTACCTCTACG | 55.55 | 2 | 72 | This study |
| | | | | | | | cdh-R | TGGAAGTGAGTTGTTCAGG | 54.63 | 0 | | This study |

| | | | | | | | | | | | | |
|---|---|---|---|---|---|---|---|---|---|---|---|---|
| Lipid metabolism | Rv2881c | **cdsA** | integral membrane phosphatidate cytidylyltransferase CdsA (CDP-diglyceride synthetase) (CDP-diglyceride pyrophosphorylase) (CDP-diacylglycerol synthase) (CDS) (CTP:phosphatidate cytidylyltransferase) (CDP-DAG synthase) (CDP-DG synthetase) | Involved in the phospholipid biosynthesis [catalytic activity: CTP + phosphatidate = pyrophosphate + CDP-diacylglycerol] | glycerophospholipid metabolism | lipid metabolism | cdsA-F | CGTTGTCTGCATGATTTGG | 55.11 | 0 | 145 | This study |
| | | | | | | | cdsA-R | GAAAGAGCAGAACAATGGG | 53.97 | 0 | | This study |
| Lipid metabolism | Rv2612c | **pgsA1** | PI synthase PgsA1 (phosphatidylinositol synthase) (CDP-diacylglycerol--inositol-3-phosphatidyltransferase) | Catalyzes the transfer of a free alcohol (inositol) onto CDP-diacylglycerol. The product of this putative ORF seems be essential to mycobacteria [catalytic activity: CDP-diacylglycerol + myo-inositol = CMP + phosphatidyl 1D-myo-inositol]. PgsA1 utilises inositol-phosphate rather than inositol, in contrast to its Eukaryotic ortholog (see Gräve et al. 2019) | glycerophospholipid metabolism | lipid metabolism | pgsA1-F | TCTGCTGTGGTGGATAGC | 56.31 | 2 | 99 | This study |
| | | | | | | | pgsA1-R | GCCTTGATGTAAGAGATCACC | 56.12 | 2 | | This study |
| Lipid metabolism | Rv0436c | **pssA** | CDP-diacylglycerol--serine O-phosphatidyltransferase PssA (PS synthase) (phosphatidylserine synthase) | Involved in phospholipid biosynthesis. Generates phosphatidylserine [catalytic activity: CDP-diacylglycerol + L-serine = CMP + O-SN-phosphatidyl-L-serine] | glycerophospholipid metabolism | lipid metabolism | pssA-F | GGGAACGTCGATACTCTTGG | 57.79 | 0 | 148 | This study |
| | | | | | | | pssA-R | CACCCAGATCAACAAGTAGGG | 58.01 | 0 | | This study |
| Lipid metabolism | Rv0437c | **psd** | phosphatidylserine decarboxylase Psd (PS decarboxylase) | Function unknown, but involved in lipid metabolism [catalytic activity: phosphatidyl-L-serine = phosphatidylethanolamine + CO(2)] | glycerophospholipid metabolism | lipid metabolism | psd-F | CACACGTCGGAGACAAGC | 56 | 2 | 83 | This study |
| | | | | | | | psd-R | CGCTGGCAGGTAGGTATCC | 56.9 | 2 | | This study |
| Lipid metabolism | Rv3229c | **desA3** | linoleoyl-CoA desaturase (delta(6)-desaturase) | Thought to be involved in lipid metabolism [catalytic activity: linoleoyl-CoA + ah(2) + O(2) = gamma-linolenoyl-CoA + a + 2 H(2)O] | linoleic acid metabolism | lipid metabolism | desA3-F | AATCGAGCATCACCTCTATCC | 57.05 | 0 | 95 | This study |
| | | | | | | | desA3-R | ATGGCAAGTCGTATCTGTCG | 57.79 | 2 | | This study |
| Carbohydrate metabolism | Rv3265c | **wbbL** | dTDP-RHA:a-D-GlcNAc-diphosphoryl polyprenol, a-3-L-rhamnosyl transferase WbbL1 (alpha-L-rhamnose-(1->3)-alpha-D-GlcNAc(1->P)-P-decaprenyl) | Probably involved in cell wall arabinogalactan linker formation: Uses dTDP-L-rhamnose as substrate to insert the rhamnosyl residue into the cell wall. Seems to be essential for mycobacterial viability | carbohydrate biosynthesis | cell wall and cell processes | wbbL-F | AGCACCTATATCTTCTTAGCC | 54.3 | 1 | 149 | This study |
| | | | | | | | wbbL-R | TCTACCAGCTTCAGTTTCC | 54.09 | 0 | | This study |
| Carbohydrate metabolism | Rv2702 | **ppgK** | Polyphosphate glucokinase PpgK (polyphosphate-glucose phosphotransferase) | Catalyzes the phosphorylation of glucose using polyphosphate or ATP as the phosphoryl donor. GTP, UTP and CTP can replace ATP as phosphoryl donor [catalytic activity: (phosphate)(N) + D-glucose = (phosphate)(N-1) + D-glucose 6-phosphate] | carbohydrate biosynthesis, glycolisis/gluconeogenesis | intermediary metabolism and respiration | ppgK-F | AAGGAAGCGGAGGAAAGG | 59.36 | 0 | 70 | This study |
| | | | | | | | ppgK-R | TGGCCCACTTTGGATAGG | 59.45 | 0 | | This study |
| Carbohydrate metabolism | Rv3667 | **acs** | Acetyl-coenzyme A synthetase Acs (acetate--CoA ligase) (acetyl-CoA synthetase) (acetyl-CoA synthase) (acyl-activating enzyme) (acetate thiokinase) (acetyl-activating enzyme) (acetate-coenzyme A ligase) (acetyl-coenzyme A synthase) | Activates acetate to acetyl-coenzyme A [catalytic activity: ATP + acetate + CoA = AMP + pyrophosphate + acetyl-CoA] | glycolysis and gluconeogenesis | intermediary metabolism and respiration | acs-F | CCTACAACTGTGTGGATCG | 55.35 | 2 | 115 | This study |
| | | | | | | | acs-R | GCAAGCAGATCGGAATAGG | 55.87 | 0 | | This study |
| Carbohydrate metabolism | Rv0761c | **adhB** | zinc-containing alcohol dehydrogenase NAD dependent AdhB | Thought to catalyze the reversible oxidation of ethanol to acetaldehyde with the concomitant reduction of NAD. [illegible] | Glycolysis/Gluconeogenesis, Fatty acid degradation, Chloroalkane and chloroalkene degradation, pyruvate, [illegible] | intermediary [illegible] | adhB-F | TGGAATGCGGAATCTGTGC | 58 | 2 | [illegible] | This study |

| | | | | | | | | | | | | |
|---|---|---|---|---|---|---|---|---|---|---|---|---|
| | | | | Probably acts on primary or secondary alcohols or hemiacetals [catalytic activity: an alcohol + NAD+ = an aldehyde or ketone + NADH] | tyrosine and methane metabolism | metabolism and respiration | adhB-R | CAGGGTCATCGGGTAGACG | 59 | 2 | 112 | This study |
| Carbohydrate metabolism | Rv3255c | **manA** | mannose-6-phosphate isomerase ManA (phosphomannose isomerase) (phosphomannoisomerase) (PMI) (phosphohexoisomerase) (phosphohexomutase) | This enzyme converts D-mannose 6-phosphate to D-fructose 6-phosphate [catalytic activity: D-mannose 6-phosphate = D-fructose 6-phosphate] | Carbohydrate biosynthesis, fructose and mannose metabolism, ManLAM biosynthesis | intermediary metabolism and respiration | manA-F | TGTTCACCACCTGGATTACC | 58.26 | 2 | 89 | This study |
| | | | | | | | manA-R | GGAGCTGACGTACTGGATAGC | 58.98 | 2 | | This study |
| Carbohydrate metabolism | Rv3264c | **manB** | D-alpha-D-mannose-1-phosphate guanylyltransferase ManB (D-alpha-D-heptose-1-phosphate guanylyltransferase) | Involved in GDP-mannose biosynthesis and biosynthesis of nucleotide-activated glycero-manno-heptose (D-alpha-D pathway): generates GDP-mannose and phosphate from GTP and alpha-D-mannose 1-phosphate. MANB product is needed for all mannosyl glycolipids and polysaccharides which, like rhamnosyl residues, are an important part of the mycobacterium envelope [catalytic activity: alpha-D-mannose 1-phosphate + GTP = GDP-mannose + phosphate] | Carbohydrate biosynthesis, fructose and mannose metabolism, ManLAM biosynthesis | cell wall and cell processes | manB-F | CTACAGATCGAATACGTGACC | 55.74 | 1 | 110 | This study |
| | | | | | | | manB-R | ACATCGCCGTTAAACACC | 55.71 | 3 | | This study |
| Carbohydrate metabolism | Rv3257c | **pmmA** | phosphomannomutase PmmA (PMM) (phosphomannose mutase) | This enzyme converses D-mannose 1-phosphate in D-mannose 6-phosphate [catalytic activity: D-mannose 1-phosphate = D-mannose 6-phosphate] | Carbohydrate biosynthesis, fructose and mannose metabolism, ManLAM biosynthesis | intermediary metabolism and respiration | pmmA-F | TTACTTCCGTGACTTCTGG | 54.15 | 0 | 121 | This study |
| | | | | | | | pmmA-R | TTCATAGCGTTGGTAGTCC | 54.31 | 1 | | This study |
| Carbohydrate metabolism | Rv3308 | **pmmB** | phosphomannomutase PmmB (phosphomannose mutase) | Converts D-mannose 1-phosphate to D-mannose 6-phosphate | Carbohydrate biosynthesis, fructose and mannose metabolism, ManLAM biosynthesis | intermediary metabolism and respiration | pmmB-F | ATACAGATCACGGCGTCACA | 59.18 | 1 | 203 | Guirado et al., 2015 |
| | | | | | | | pmmB-R | CGCTGGATATAACGGTCGAT | 57.37 | 2 | | Guirado et al., 2015 |
| Carbohydrate metabolism | Rv3631 | **Rv3631** | Possible transferase (possibly glycosyltransferase) | Function unknown; probably involved in cellular metabolism | ManLAM biosynthesis | intermediary metabolism and respiration | Rv3631-F | CTTGACCGACACCAACAATG | 57.31 | 3 | 224 | Guirado et al., 2015 |
| | | | | | | | Rv3631-R | GAAACCCGTCGAAAATGATG | 55.38 | 1 | | Guirado et al., 2015 |
| Carbohydrate metabolism | Rv2051c | **ppm1** | Polyprenol-monophosphomannose synthase Ppm1 | Transfers mannose from GDP-mannose to all endogenous polyprenol-phosphates | carbohydrate biosynthesis, ManLAM biosynthesis | cell wall and cell processes | ppm1-F | CTCCAAGGGCTACTGCTTCC | 60.11 | 0 | 93 | This study |
| | | | | | | | ppm1-R | CGCTCGGTAAAGGTAATCG | 55.95 | 2 | | This study |
| Carbohydrate metabolism | Rv3260c | **whiB2** | transcriptional regulatory protein WhiB-like WhiB2 | Involved in transcriptional mechanism | Transcription factor, redox homeostasis | regulatory proteins | whiB2-F | GCACGAGTGTCTGGAGTACG | 57.6 | 2 | 99 | This study |
| | | | | | | | whiB2-R | ATGATCCCGCGTTTGAGG | 54.9 | 0 | | This study |
| Carbohydrate metabolism | Rv3441c | **mrsA** | phospho-sugar mutase / MrsA protein homolog | Function unknown; involved in cellular metabolism | carbohydrate biosynthesis, mannose donor biosynthesis pathway, Amino sugar and nucleotide sugar metabolism, UDP-glucose biosynthesis | intermediary metabolism and respiration | mrsA-F | CGCTGGCAATGAAAGAGG | 56.49 | 1 | 71 | This study |
| | | | | | | | mrsA-R | CCCGAGGTTACTCATCACG | 57.36 | 3 | | This study |
| Carbohydrate metabolism | Rv3068c | **pgmA** | phosphoglucomutase PgmA | This enzyme participates in both the breakdown and synthesis of glucose [catalytic activity: alpha-D-glucose 1-phosphate = alpha-D-glucose 6-phosphate] | Carbohydrate metabolism: carbohydrate biosynthesis, mannose donor biosynthesis pathway, Galactose metabolism, Glycolysis/ gluconeogenesis, penthose phosphate pathway, amino and nucleotide sugar metabolism | intermediary metabolism and respiration | pgmA-F | GGTCGGATTCAAATGGTTCG | 57.17 | 2 | 90 | This study |
| | | | | | | | pgmA-R | CGTCGCAGAAATGATGCC | 57.02 | 1 | | This study |
| Carbohydrate metabolism | Rv0363c | **fba** | fructose-bisphosphate aldolase Fba | Involved in glycolysis (at the sixth step) [catalytic activity: D-fructose 1,6-bisphosphate = glycerone phosphate + D-glyceraldehyde 3-phosphate] | fructose and mannose metabolism | intermediary metabolism and respiration | fba-F | GGTGTCAAGAAGGTCTACG | 54.98 | 2 | 117 | This study |
| | | | | | | | fba-R | GTGGGTTAGGGACTTTCC | 54.98 | 3 | | This study |
| Carbohydrate metabolism | Rv0727c | **fucA** | L-fuculose phosphate aldolase FucA (L-fuculose-1-phosphate aldolase) | Involved in fucose metabolism (at the third step) [catalytic activity: L-fuculose 1-phosphate = glycerone phosphate + (S)-lactaldehyde] | fructose and mannose metabolism | intermediary metabolism and respiration | fucA-F | GGAAATATCTCAGCCAGGCG | 58.48 | 3 | 84 | This study |
| | | | | | | | fucA-R | ATCGTGGAGCAGCATCTCG | 59.93 | 3 | | This study |
| Carbohydrate metabolism | Rv3010c | **pfkA** | Probable 6-phosphofructokinase PfkA (phosphohexokinase) (phosphofructokinase) | Involved in glycolysis; converts sugar-1-P to sugar-1,6-P [catalytic activity: ATP + D-fructose 6-phosphate = ADP + D-fructose 1,6-bisphosphate]. | Fructose and mannose, Galactose, Glycolysis/Gluconeogenesis, and penthose metabolism | intermediary metabolism and respiration | pfkA-F | TCACATGACCCTGATTCC | 53.44 | 1 | 96 | This study |
| | | | | | | | pfkA-R | CAGATGAAATGCGAGTCC | 53.24 | 1 | | This study |

| | | | | | | | | | | | | |
|---|---|---|---|---|---|---|---|---|---|---|---|---|
| Carbohydrate metabolism | Rv1099c | **glpX** | Fructose 1,6-bisphosphatase GlpX | Involved in gluconeogenesis [catalytic activity: fructose-1,6-bisphosphate + H2O = D-fructose-6-phosphate + phosphate] | glycolysis/ gluconeogenesis, fructose and mannose metabolism, penthose phosphate pathway | intermediary metabolism and respiration | glpX-F | TCAAAGCTCAACGAATACTCC | 56 | 2 | 68 | This study |
| | | | | | | | glpX-R | TAGGGCAATGGGTACACG | 56.3 | 2 | | This study |
| Carbohydrate metabolism | Rv0620 | **galK** | galactokinase GalK (galactose kinase) | Involved in galactose metabolism (leloir pathway) (at the first reaction) [catalytic activity: ATP + D-galactose = ADP + D-galactose 1-phosphate] | galactose metabolism | intermediary metabolism and respiration | galK-F | CTGACCGAGAATCAGCGGG | 60.23 | 1 | 61 | This study |
| | | | | | | | galK-R | CGGTGAAATCCGAATCAGCC | 59.34 | 3 | | This study |
| Carbohydrate metabolism | Rv0618 | **galT** | galactose-1-phosphate uridylyltransferase GalTa [first part] | Involved in galactose metabolism (leloir pathway) [catalytic activity: UTP + alpha-D-galactose 1-phosphate = diphosphate + UDP-galactose] | galactose metabolism | intermediary metabolism and respiration | galT-F | TGATCGGATGTATTCTTCACC | 55.21 | 1 | 133 | This study |
| | | | | | | | galT-R | CGCCAGATATTTCAGTTTGG | 54.76 | 0 | | This study |
| Carbohydrate metabolism | Rv0993 | **galU** | UTP--glucose-1-phosphate uridylyltransferase GalU (UDP-glucose pyrophosphorylase) (UDPGP) (alpha-D-glucosyl-1-phosphate uridylyltransferase) (uridine diphosphoglucose pyrophosphorylase) | May play a role in stationary phase survival [catalytic activity: UTP + alpha-D-glucose 1-phosphate = diphosphate + UDP-glucose] | galactose metabolism, pentose and glucuronate conversion | intermediary metabolism and respiration | galU-F | CGCATTTCGTGGAAGACC | 56.54 | 0 | 106 | This study |
| | | | | | | | galU-R | GATTCGACCTTGATCAGTGC | 56.62 | 2 | | This study |
| Carbohydrate metabolism | Rv2471 | **aglA** | alpha-glucosidase AglA (maltase) (glucoinvertase) (glucosidosucrase) (maltase-glucoamylase) (lysosomal alpha-glucosidase) (acid maltase) | Involved in sugar metabolism (hydrolysis of terminal, non-reducing 1,4-linked D-glucose residues with release of D-glucose) | galactose metabolism | intermediary metabolism and respiration | aglA-F | CAATCACGATGTGGGACGG | 58.62 | 1 | 127 | This study |
| | | | | | | | aglA-R | TTCCTGGCCGTTGTAGAGG | 59.02 | 0 | | This study |
| Carbohydrate metabolism | Rv0211 | **pckA** | iron-regulated phosphoenolpyruvate carboxykinase [GTP] PckA (phosphoenolpyruvate carboxylase) (PEPCK)(pep carboxykinase) | Rate-limiting gluconeogenic enzyme [catalytic activity: GTP + oxaloacetate = GDP + phosphoenolpyruvate + CO2] | citrate cycle, pyruvate metabolism | intermediary metabolism and respiration | pckA-F | CCGAGAAGCACAAGAACTCC | 58.57 | 1 | 206 | Guirado et al., 2015 |
| | | | | | | | pckA-R | ACAGAACGGCACCACATACA | 59.61 | 0 | | Guirado et al., 2015 |
| Carbohydrate metabolism | Rv0462 | **lpdC** | Dihydrolipoamide dehydrogenase LpdC (lipoamide reductase (NADH)) (lipoyl dehydrogenase) (dihydrolipoyl dehydrogenase) (diaphorase) | Involved in energy metabolism. Lipoamide dehydrogenase is the E3 component of pyruvate dehydrogenase and the alpha-ketoacid dehydrogenase complex [catalytic activity: dihydrolipoamide + NAD(+) = lipoamide + NADH]. Also involved in antioxidant defense; LPDC\|Rv0462, DLAT\|Rv2215, AHPD\|Rv2429, and AHPC\|Rv2428 constitute an NADH-dependent peroxidase and peroxynitrite reductase that provides protection against oxidative stress. | citrate cycle, Glycolysis/Gluconeogenesis | intermediary metabolism and respiration | lpdC-F | CCTCAATGTCGGCTGTATCC | 55.2 | 2 | 74 | This study |
| | | | | | | | lpdC-R | CGTCCTTGGTGAAGATGTGG | 55.7 | 0 | | This study |
| Carbohydrate metabolism | Rv0794c | **Rv0794c** | oxidoreductase | Function unknown; probably involved in cellular metabolism | citrate cycle, Glycine, serine and threonine metabolism, Glycolysis / Gluconeogenesis, Glyoxylate and dicarboxylate metabolism, pyruvate metabolism | intermediary metabolism and respiration | Rv0794c-F | CCCACCAAGGCAAATACC | 55.59 | 2 | 145 | This study |
| | | | | | | | Rv0794c-R | GGGTCGGTAAAGAATGCC | 55.42 | 1 | | This study |
| Carbohydrate metabolism | Rv3075c | **Rv3075c** | Citrate lyase beta chain | Unknown | citrate cycle | conserved hypotheticals | Rv3075c-F | GAAATCGTCTGGGCGAAGG | 58.62 | 0 | 190 | This study |
| | | | | | | | Rv3075c-R | AGTGGTAGGTGTCAGTCGG | 58.35 | 1 | | This study |
| Carbohydrate metabolism | Rv0248c | **Rv0248c** | Probable succinate dehydrogenase [iron-sulfur subunit] (succinic dehydrogenase) | Involved in interconversion of fumarate and succinate (aerobic respiration) [catalytic activity: succinate + acceptor = fumarate + reduced acceptor] | Butanoate and carbon metabolism, citrate cycle | intermediary metabolism and respiration | Rv0248c-F | AGTGTCAAGGGAATTCTGG | 54.34 | 0 | 94 | This study |
| | | | | | | | Rv0248c-R | GAATGTAGTCGAACATGAAGC | 55.08 | 2 | | This study |
| Carbohydrate metabolism | Rv3339c | **icd1** | isocitrate dehydrogenase [NADP] Icd1 (oxalosuccinate decarboxylase) (IDH) (NADP+-specific ICDH) (IDP) | Involved in the KREBS cycle [catalytic activity: isocitrate + NADP(+) = 2-oxoglutarate + CO(2) + NADPH] | cytrate cycle, 2-Oxocarboxylic acid cycle, glutathione metabolism | intermediary metabolism and respiration | icd1-F | ATCTGTCCACCAAGAACACC | 57.38 | 0 | 60 | This study |
| | | | | | | | icd1-R | CGAACTCGTCTTTGAACATCC | 59.73 | 0 | | This study |

| | | | | | | | | | | | | |
|---|---|---|---|---|---|---|---|---|---|---|---|---|
| Carbohydrate metabolism | Rv3316 | **sdhC** | succinate dehydrogenase (cytochrome B-556 subunit) SdhC (succinic dehydrogenase) (fumarate reductase) (fumarate dehydrogenase) (fumaric hydrogenase) | Involved in tricarboxylic acid cycle. Mono-heme cytochrome of the succinate dehydrogenase complex | cytrate cycle, Butanoate metabolism | intermediary metabolism and respiration | sdhC-F | TCCTGTTTGTCCATGTCC | 54.04 | 2 | 163 | This study |
| | | | | | | | sdhC-R | ATCAAGATGACCCGAATCC | 54.31 | 2 | | This study |
| Carbohydrate metabolism | Rv1915 | **aceA** | isocitrate lyase AceAa [first part] (isocitrase) (isocitratase) (Icl) | Involved in glyoxylate bypass, an alternative to the tricarboxylic acid cycle [catalytic activity: isocitrate = succinate + glyoxylate] | glyoxylate and dicarboxylate metabolism | intermediary metabolism and respiration | aceA-F | ACGTACCGTCCTACAAGTCC | 56.64 | 0 | 94 | This study |
| | | | | | | | aceA-R | AAGCGCATAGAGAAGATGACC | 58.57 | 1 | | This study |
| Carbohydrate metabolism | Rv0467 | **icl1** | Isocitrate lyase Icl (isocitrase) (isocitratase) | Involved in glyoxylate bypass (at the first step), an alternative to the tricarboxylic acid cycle (in bacteria, plants, and fungi) [catalytic activity: isocitrate = succinate + glyoxylate]. Involved in the persistence in the host | glyoxylate and dicarboxylate metabolism | intermediary metabolism and respiration | icl1-F | TCAAGTTCCAGTTCATCACG | 55.71 | 2 | 111 | This study |
| | | | | | | | icl1-R | CCTGCAGTTCGACATACG | 55.63 | 2 | | This study |
| Carbohydrate metabolism | Rv0046c | **ino1** | myo-inositol-1-phosphate synthase Ino1 (inositol 1-phosphate synthetase) (D-glucose 6-phosphate cycloaldolase) (glucose 6-phosphate cyclase) (glucocycloaldolase) | Involved in phosphatidylinositol (PI) biosynthetic pathway [catalytic activity: D-glucose 6-phosphate = 1L-myo-inositol 1-phosphate] | inositol phosphate metabolism | intermediary metabolism and respiration | ino1-F | ATGGACTTCCTCAACATGC | 55.16 | 2 | 60 | This study |
| | | | | | | | ino1-R | GGTCTTGGAGATCTTCTTGG | 55.15 | 2 | | This study |
| Others | Rv3423c | **alr** | Alanine racemase Alr | Provides the D-alanine required for cell wall biosynthesis. Transforms L-alanine to D-alanine [catalytic activity: L-alanine = D-alanine] | Amino acid metabolism: Alanine metabolism, Vancomycin resistance | intermediary metabolism and respiration | alr-F | AAACGACACGACAGATGG | 55.02 | 0 | 116 | This study |
| | | | | | | | alr-R | GCACGTTGTGTTCAATAGC | 55.46 | 2 | | This study |
| Others | Rv0391 | **metZ** | O-succinylhomoserine sulfhydrylase MetZ (OSH sulfhydrylase) | Involved in methionine biosynthesis. Converts O-succinylhomoserine into homocysteine | Amino acid metabolism: Cysteine and methionine metabolism | intermediary metabolism and respiration | metZ-F | TATGTGTACTCCCGCTACG | 56 | 2 | 185 | This study |
| | | | | | | | metZ-R | AAACACGAGCCAAACAGG | 55.82 | 0 | | This study |
| Others | Rv1295 | **thrC** | Threonine synthase ThrC (ts) | Involved in threonine biosynthesis [catalytic activity: O-phospho-L-homoserine + H(2)O = L-threonine + phosphate] | Amino acid metabolism: Glycine, serine and threonine metabolism, Vitamin B6 metabolism | intermediary metabolism and respiration | thrC-F | GGCAACTAATCTCTCCAAGC | 56.2 | 2 | 96 | This study |
| | | | | | | | thrC-R | ATCGTCATGCCACGATCC | 57.24 | 2 | | This study |
| Others | Rv1381 | **pyrC** | dihydroorotase PyrC (DHOase) | Involved in pyrimidine biosynthesis (third step) [catalytic activity: (S)-dihydroorotate + H(2)O = N-carbamoyl-L-aspartate] | Amino acid metabolism: Pyrimidine metabolism | intermediary metabolism and respiration | pyrC-F | GTGCTGATTCGTGGTGTGC | 59 | 2 | 91 | This study |
| | | | | | | | pyrC-R | GATCCGGTCCTATCTGGGC | 59 | 2 | | This study |
| Others | Rv1601 | **hisB** | imidazole glycerol-phosphate dehydratase HisB | Histidine biosynthesis (seventh step) [catalytic activity: D-erythro-1-(imidazol-4-YL)glycerol 3- phosphate = 3-(imidazol-4-YL)-2-oxopropyl phosphate + H(2)O] | Amino acid metabolism: Histidine metabolism | intermediary metabolism and respiration | hisB-F | CGATGGACGAAACACTGG | 55.83 | 1 | 66 | This study |
| | | | | | | | hisB-R | CGGTATGCACGCAATAGG | 56.07 | 0 | | This study |
| Others | Rv1603 | **hisA** | phosphoribosylformimino-5-aminoimidazole carboxamide ribotide isomerase HisA | Histidine biosynthesis pathway (fourth step) [catalytic activity: N-(5'-phospho-D-ribosylformimino)-5-amino-1-(5''-phosphoribosyl)-4-imidazolecarb oxamide = N-(5'-phospho-D-1'-ribulosylformimino)-5-amino-1-(5''-phosphoribosyl)-4-imidazolecarboxamide.] | Amino acid metabolism: Histidine metabolism, Phenylalanine, tyrosine and tryptophan biosynthesis | intermediary metabolism and respiration | hisA-F | TAGACAGTGAAGGATGTTCGC | 57.77 | 2 | 83 | This study |
| | | | | | | | hisA-R | CAGCAGGTCCAGATTGGG | 57.35 | 2 | | This study |
| Others | Rv1655 | **argD** | acetylornithine aminotransferase ArgD | Arginine biosynthesis (fourth step) [catalytic activity: N2-acetyl-L-ornithine + 2-oxoglutarate = N-acetyl-L-glutamate 5-semialdehyde + L-glutamate] | Amino acid metabolism: 2-Oxocarboxylic acid metabolism, Arginine biosynthesis, Lysine biosynthesis | intermediary metabolism and respiration | argD-F | GTGATGATGAACAACTACGG | 54.16 | 1 | 100 | This study |
| | | | | | | | argD-R | CGAGCAGGTCGATATAGG | 53.77 | 2 | | This study |
| Others | Rv2589 | **gabT** | 4-aminobutyrate aminotransferase GabT (gamma-amino-N-butyrate transaminase) (GABA transaminase) | Involved in 4-aminobutyrate (GABA) degradation pathway [catalytic activity: 4-aminobutanoate + 2-oxoglutarate = succinate semialdehyde + L-glutamate] | Amino acid metabolism: GABA degradation, Alanine, aspartate and glutamate metabolism, Butanoate metabolism, Propanoate metabolism, Valine, leucine and isoleucine | intermediary metabolism and | gabT-F | CGAGCGCCATTGTCTTACC | 58.99 | 0 | 171 | This study |

| | | | | | | | | | | | | |
|---|---|---|---|---|---|---|---|---|---|---|---|---|
| | | | transaminase) (glutamate:succinic semialdehyde transaminase) (GABA aminotransferase) (GABA-at) | | Valine, leucine and isoleucine degradation | metabolism and respiration | gabT-R | AACGATGAAACCGCCTTCG | 58.99 | 2 | 171 | This study |
| Others | Rv0884c | **serC** | phosphoserine aminotransferase SerC (PSAT) | Catalyzes the reversible interconversion of phosphoserine and 2-oxoglutarate to 3-phosphonooxypyruvate and glutamate. Require both in the major phosphorylated pathway of serine biosynthesis and in pyridoxine biosynthesis [catalytic activity: O-phospho-L-serine + 2-oxoglutarate = 3-phosphonooxypyruvate + L-glutamate] | Carbohydrate metabolism/ Amino acid metabolism: Glycine, serine and threonine metabolism, Methane metabolism, Vitamin B6 metabolism, biosynthesis of amino acids | intermediary metabolism and respiration | serC-F | GCTGGGTTCCTGATTTCC | 55.29 | 0 | 74 | This study |
| | | | | | | | serC-R | CGGTGTGTTGTATGTCTGG | 55.96 | 0 | | This study |
| Others | Rv0283 | **eccB3** | ESX conserved component EccB3. ESX-3 type VII secretion system protein. Possible membrane protein | Unknown | ESX-3 secretion system | cell wall and cell processes | eccB3-F | GGCTGTTCAACGCGATACC | 59.29 | 2 | 136 | This study |
| | | | | | | | eccB3-R | TTGCCCGAGTCTTTCAGGG | 59.63 | 1 | | This study |
| Others | Rv1984c | **cfp21** | cutinase precursor CFP21 | Hydrolyzes cutin. Shown to have esterase and lipase activity | Not assigned | cell wall and cell processes | cfp21-F | AAGACCATAAACTTGTGTGC | 54.11 | 2 | 82 | This study |
| | | | | | | | cfp21-R | ACTGAACATACGAAACATGC | 54.34 | 2 | | This study |
| Others | Rv2428 | **ahpC** | Alkyl hydroperoxide reductase C protein AhpC (alkyl hydroperoxidase C) | Involved in oxidative stress response. LPDC\|Rv0462, DLAT\|Rv2215, AHPD\|Rv2429, and AHPC\|Rv2428 constitute an NADH-dependent peroxidase and peroxynitrite reductase that provides protection against oxidative stress. Response of *Mtb* to phagocytosis, tolerance by *Mtb* to nitric oxide produced by macrophages | Redox homeostasis | virulence, detoxification, adaptation | ahpC-F | CGCGTGACCTTTATCGTCG | 59.09 | 3 | 185 | This study |
| | | | | | | | ahpC-R | GAAGCCTTGAGGAGTTCGC | 58.54 | 2 | | This study |
| Others | Rv0432 | **sodC** | Periplasmic superoxide dismutase [Cu-Zn] SodC | Destroys radicals which are normally produced within the cells and are toxic to biological systems [catalytic activity: 2 superoxide + 2 H+ = O2 + H2O2] | Redox homeostasis | virulence, detoxification, adaptation | sodC-F | ACTTTGCCAACATTCCGCC | 59.33 | 1 | 118 | This study |
| | | | | | | | sodC-R | GAACCAATGACACCGCACG | 59.79 | 2 | | This study |
| Others | Rv3082c | **virS** | Virulence-regulating transcriptional regulator VirS (AraC/XylS family) | May have a role in the regulation of proteins necessary for virulence | Not assigned | virulence, detoxification, adaptation | virS-F | CTACCTCTACGTCCATTCG | 54.26 | 2 | 92 | This study |
| | | | | | | | virS-R | GTTCGGTCACCTCATAGC | 54.85 | 2 | | This study |
| Others | Rv3133c | **dosR** | Two component transcriptional regulatory protein DevR (probably LuxR/UhpA-family) | Regulator part of the two component regulatory system DEVR/DEVS/dost. Controls HSPX\|Rv2031\|ACR expression | Two-component system | regulatory proteins | dosR-F | GTCTGGTTGACTTGCTTGGG | 59.05 | 0 | 150 | This study |
| | | | | | | | dosR-R | ACAGTTCAATGCCGTTGCC | 59.34 | 1 | | This study |
| Others | Rv0757 | **PhoP** | two component system response transcriptional positive regulator PhoP | Involved in transcriptional mechanism. Part of the two component regulatory system PHOP/PHOQ. This protein is thought to be a positive regulator for the phosphate regulon, required for intracellular growth. Transcription of this operon is positively regulated by PHOB and PHOR\|Rv0758 when phosphate is limited | Two-component system | regulatory proteins | phoP-F | GACAAAGCCCTTCAGTTTGG | 54.3 | 0 | 88 | This study |
| | | | | | | | phoP-R | ATTACGTGGTTCCTTGTTGC | 53.7 | 2 | | This study |

**Supplemental Table 2**

| Category | Gene | A1 | A2 | A3 | E1 | E2 | E3 |
|---|---|---|---|---|---|---|---|
| Lipid metabolism | **pimA** | 7.917490773 | 5.275966773 | 2.321286604 | 1.22518783 | 1.019394192 | 1.995417477 |
| Lipid metabolism | **pimF** | 23.69407391 | 3.321308829 | 1.59309755 | 0.847128571 | 1.263769835 | 1.056228661 |
| Lipid metabolism | **pgsA** | 14.85552866 | 5.510552648 | 2.477255798 | 0.925409412 | 0.859374223 | 2.06750063 |
| Lipid metabolism | **mgtA** | 10.2459103 | 3.673707172 | 2.412506476 | 1.235009452 | 1.165058382 | 1.123963807 |
| Lipid metabolism | **embC** | 12.76512837 | 5.625237364 | 2.125660244 | 1.469160842 | 1.171555254 | 2.313780309 |
| Lipid metabolism | **embR** | 198.3041959 | 7.009884625 | 3.802249975 | 1.030335633 | 1.25717439 | 1.059879261 |
| Lipid metabolism | **Rv1747** | 0.111620819 | 0.145827987 | 0.093851872 | 0.529124486 | 1.542713485 | 2.913430062 |
| Lipid metabolism | **Rv2623** | 62.53807823 | 3.769666796 | 3.574891259 | 0.868714975 | 1.041429741 | 0.869600641 |
| Lipid metabolism | **acpM** | 183.1827795 | 5.293989486 | 3.596401979 | 1.210600611 | 1.21881887 | 1.322283451 |
| Lipid metabolism | **kasA** | 168.6971255 | 5.105935987 | 2.676398821 | 1.085481995 | 1.101355305 | 1.609087197 |
| Lipid metabolism | **kasB** | 9.703545341 | 3.66238075 | 1.785387738 | 0.791293645 | 1.318685186 | 1.161943544 |
| Lipid metabolism | **pks13** | 32.68773318 | 6.784665425 | 3.422551255 | 0.752246737 | 0.81318967 | 2.321524469 |
| Lipid metabolism | **inhA** | 158.2631849 | 6.554835259 | 3.308255814 | 1.04720214 | 1.215264276 | 1.830937496 |
| Lipid metabolism | **fabH** | 0.305227717 | 0.306091705 | 0.309266405 | 1.226327036 | 1.115341577 | 1.505500021 |
| Lipid metabolism | **fabG4** | 14.19057781 | 10.33319894 | 3.688714621 | 0.106582933 | 0.483450634 | 2.3840906 |
| Lipid metabolism | **fadA2** | 127.6776181 | 5.068971576 | 4.500431628 | 0.753104216 | 1.103799894 | 0.97198206 |
| Lipid metabolism | **fadD25** | 0.481307794 | 0.528024684 | 0.497056036 | 0.820857793 | 1.37362134 | 1.007165958 |
| Lipid metabolism | **echA3** | 131.6652634 | 6.232047862 | 3.277243606 | 0.902135682 | 1.106388476 | 0.962969243 |
| Lipid metabolism | **alkB** | 50.44510122 | 4.306742223 | 3.178214854 | 0.394484505 | 0.907488407 | 1.438747021 |
| Lipid metabolism | **fadB** | 103.6996755 | 5.011291257 | 3.884676137 | 0.80704103 | 1.119529192 | 0.752065849 |
| Lipid metabolism | **fadE5** | 26.55671608 | 5.764677223 | 5.056542949 | 0.598204353 | 1.017142606 | 1.268078697 |
| Lipid metabolism | **adhE1** | 0.151808498 | 0.438746508 | 0.472796101 | 1.008898497 | 1.047421923 | 1.410437001 |
| Lipid metabolism | **glpK** | 0.065978112 | 0.137098173 | 0.183857809 | 0.066155339 | 0.932661709 | 1.701387045 |
| Lipid metabolism | **lipY** | 51.55230971 | 6.463475756 | 2.681297704 | 1.094280535 | 0.927258696 | 1.029414681 |
| Lipid metabolism | **tgs1** | 81.54565551 | 3.814245106 | 4.061711292 | 0.631561104 | 0.900662139 | 1.022309822 |
| Lipid metabolism | **plsB1** | 14.68339828 | 2.278035874 | 1.18451348 | 1.039684695 | 1.335814165 | 1.426441756 |
| Lipid metabolism | **gpdA1** | 8.838695654 | 2.917283464 | 1.841126086 | 0.794871457 | 0.906129795 | 1.270835273 |
| Lipid metabolism | **cdh** | 6.799478534 | 2.783312233 | 2.102071194 | 0.946438243 | 0.860371958 | 1.227952441 |
| Lipid metabolism | **cdsA** | 0.053693079 | 0.208413863 | 0.21039275 | 0.79618777 | 1.032469286 | 1.649325016 |
| Lipid metabolism | **pgsA1** | 13.05791985 | 5.287410429 | 3.250782784 | 0.851936531 | 0.766804561 | 1.764392046 |
| Lipid metabolism | **pssA** | 56.95807208 | 11.38196593 | 3.342939625 | 0.581867473 | 1.114402569 | 2.805978249 |
| Lipid metabolism | **psd** | 54.54497566 | 5.789110458 | 3.502932966 | 1.136933134 | 0.998844487 | 1.219619262 |
| Lipid metabolism | **desA3** | 103.2838756 | 5.075014488 | 5.140939118 | 0.784872108 | 0.826325133 | 0.841189462 |

| | | | | | | | |
|---|---|---|---|---|---|---|---|
| Carbohydrate metabolism | **wbbL** | 0.252289869 | 0.327708011 | 0.160272423 | 0.04153904 | 1.365637036 | 2.093383458 |
| Carbohydrate metabolism | **ppgK** | 0.377265352 | 0.427002478 | 0.577711281 | 0.733722649 | 1.169348369 | 0.818044302 |
| Carbohydrate metabolism | **acs** | 1.867207867 | 2.536859252 | 1.700574061 | 0.716206692 | 0.798408913 | 1.787017864 |
| Carbohydrate metabolism | **adhB** | 120.977342 | 5.650369887 | 3.397492562 | 0.727512002 | 1.113888286 | 1.16284614 |
| Carbohydrate metabolism | **manA** | 59.13142223 | 6.17886225 | 3.964878195 | 0.930508102 | 0.773191568 | 1.197111431 |
| Carbohydrate metabolism | **manB** | 60.9252486 | 5.652986968 | 4.146679949 | 1.207170684 | 0.948300125 | 1.37180688 |
| Carbohydrate metabolism | **pmmA** | 0.026211216 | 0.157886167 | 0.203309053 | 0.611590118 | 1.267964862 | 0.781854522 |
| Carbohydrate metabolism | **Rv3631** | 14.27989838 | 8.447557311 | 3.179416009 | 0.898486407 | 1.079273525 | 1.235322863 |
| Carbohydrate metabolism | **ppm1** | 3.179378553 | 1.076847705 | 0.757280796 | 1.127750373 | 1.833258796 | 0.708552249 |
| Carbohydrate metabolism | **whiB2** | 100.7232407 | 5.314678816 | 4.498859403 | 0.773454732 | 1.122757334 | 0.851316027 |
| Carbohydrate metabolism | **mrsA** | 71.09754626 | 5.309449903 | 3.208446815 | 0.807816601 | 1.106522062 | 1.264777581 |
| Carbohydrate metabolism | **pgmA** | 182.6555589 | 6.5726993 | 4.402192368 | 0.966432355 | 1.079988902 | 1.002855224 |
| Carbohydrate metabolism | **fba** | 2.145096668 | 0.355235704 | 0.681881901 | 0.801437354 | 0.75703831 | 0.764883536 |
| Carbohydrate metabolism | **fucA** | 170.4807489 | 7.0853746 | 5.477149626 | 0.178152954 | 1.080452754 | 0.807545196 |
| Carbohydrate metabolism | **pfkA** | 0.0878992 | 0.054293779 | 0.071180254 | 0.624495156 | 1.774913624 | 1.098066923 |
| Carbohydrate metabolism | **glpX** | 28.00249814 | 3.830907106 | 2.835509879 | 0.988027097 | 1.168452681 | 0.902832602 |
| Carbohydrate metabolism | **galK** | 144.6064727 | 7.803913416 | 3.618703327 | 1.265465404 | 0.882589052 | 0.701586658 |
| Carbohydrate metabolism | **galT** | 1.693612182 | 1.618895788 | 0.743502846 | 1.220311839 | 0.720138436 | 1.501042748 |
| Carbohydrate metabolism | **galU** | 97.00112493 | 9.672755895 | 4.804026711 | 0.886664838 | 0.488314837 | 1.742361701 |
| Carbohydrate metabolism | **aglA** | 3.566998173 | 11.05930443 | 3.681619129 | 0.710094873 | 0.789297108 | 5.11021543 |
| Carbohydrate metabolism | **pckA** | 15.52508775 | 3.341934802 | 3.969670145 | 0.37072449 | 0.980729262 | 0.521756121 |
| Carbohydrate metabolism | **lpdC** | 95.92944011 | 4.459871521 | 2.422463499 | 1.125819328 | 1.415844226 | 1.174435327 |
| Carbohydrate metabolism | **Rv3075c** | 1.49534119 | 5.740365503 | 3.913868622 | 0.452680037 | 0.870143812 | 1.34739587 |
| Carbohydrate metabolism | **Rv0248c** | 2.137918514 | 1.109180539 | 0.91199724 | 0.864999508 | 1.201454676 | 1.241105653 |
| Carbohydrate metabolism | **icd1** | 13.07682271 | 1.063442209 | 0.745738779 | 1.134837592 | 1.239349606 | 1.189925418 |
| Carbohydrate metabolism | **sdhC** | 0.010996898 | 0.044748187 | 0.087396309 | 0.540358866 | 1.146913126 | 0.868817857 |
| Carbohydrate metabolism | **aceA** | 139.496883 | 4.665123666 | 2.843414478 | 0.670663026 | 1.097874898 | 0.831162712 |
| Carbohydrate metabolism | **icl1** | 50.38517245 | 2.826421125 | 4.14295998 | 0.465572393 | 0.734005501 | 0.469117089 |
| Carbohydrate metabolism | **ino1** | 1.338528789 | 0.479602475 | 0.49565418 | 0.797248389 | 1.811510524 | 0.59286105 |
| Others | **alr** | 31.89148731 | 7.373677283 | 5.279337497 | 0.760366947 | 0.954631902 | 0.711599953 |
| Others | **thrC** | 127.7320637 | 7.170195649 | 3.67019896 | 1.142101869 | 0.971539252 | 1.764441714 |
| Others | **pyrC** | 164.1687062 | 9.082583854 | 3.684164874 | 1.342993386 | 1.315426525 | 1.331859287 |
| Others | **hisA** | 95.48428596 | 6.327534697 | 4.043094097 | 1.123235221 | 1.018482155 | 1.855817438 |
| Others | **hisB** | 78.29179701 | 6.052608844 | 4.104884687 | 0.724812704 | 1.016434212 | 0.909068532 |
| Others | **argD** | 3.849805017 | 0.81143308 | 2.669270192 | 0.38922385 | 0.381572036 | 0.626166749 |
| Others | **gabT** | 82.07240883 | 8.708627351 | 4.039519613 | 1.069951459 | 0.903732654 | 1.525653189 |

| | | | | | | | |
|---|---|---|---|---|---|---|---|
| Others | **serC** | 0.063735562 | 0.068111141 | 0.11453414 | 0.779954537 | 1.246002888 | 1.050841677 |
| Others | **eccB3** | 312.9301191 | 10.64372597 | 7.17789879 | 1.063360794 | 0.466061998 | 1.240018745 |
| Others | **cfp21** | 2.37951323 | 1.297018987 | 1.013657518 | 0.946835116 | 1.093686172 | 1.028293745 |
| Others | **ahpC** | 78.04289075 | 8.312189853 | 3.030271884 | 1.139280145 | 1.09559897 | 1.271475325 |
| Others | **sodC** | 121.7999 | 8.598213728 | 4.274247873 | 1.170709394 | 1.021269011 | 1.573007097 |
| Others | **virS** | 1.606999948 | 0.495330104 | 1.23858917 | 0.849755523 | 1.008941921 | 0.803758582 |
| Others | **dosR** | 17.07510104 | 4.234638857 | 2.619709924 | 0.588859466 | 0.922500863 | 1.457801259 |
| Others | **phoP** | 62.60195166 | 4.155473819 | 3.484535942 | 0.811432997 | 1.048783267 | 1.075243488 |

**Supplemental Fig 1**

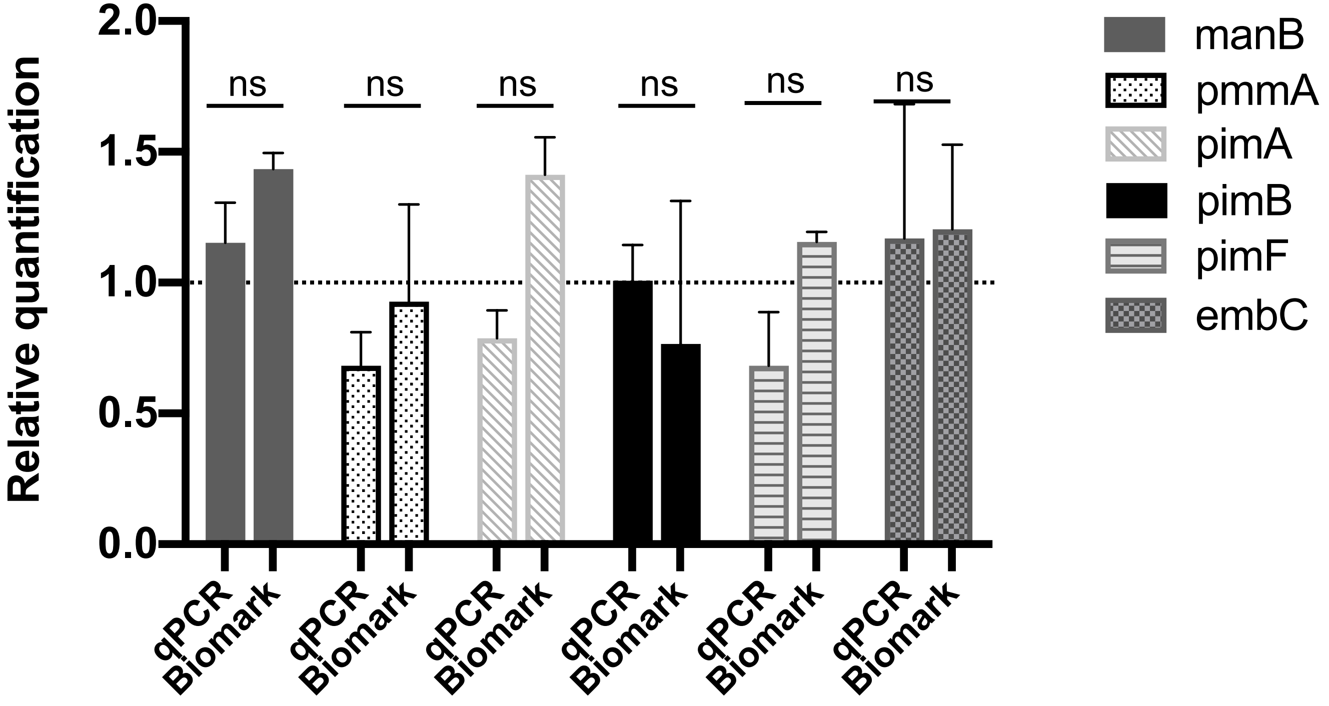
